# Enzyme-Constrained Metabolic Modeling of the Tripartite Synapse Links Astrocyte-Neuron Lactate Shuttle to Lipid Trafficking and Redox Homeostasis

**DOI:** 10.64898/2026.06.15.732277

**Authors:** Samuel Hayford Ayensu, Dan Hideki Horimoto, Carlos C. Flores, Jason R. Gerstner, Wheaton L. Schroeder

## Abstract

Neuron–astrocyte metabolic coupling sustains synaptic function through coordinated substrate exchange, redox homeostasis, and lipid handling, yet the systems-level mechanism remains poorly understood. Here, we developed an enzyme-constrained, genome-scale metabolic model of the mouse tripartite synapse to test whether protein allocation can reproduce the astrocyte– neuron lactate shuttle (ANLS). Starting from the iMM1865 mouse metabolic model, we corrected mass and charge imbalances, expanded fatty acid and lipid pathways, and standardized annotations to obtain a high-quality, generic mouse model, improving the MEMOTE score from 46% to 83%. Cell type–specific astrocyte and neuron models were reconstructed using the integrative metabolic analysis tool (iMAT) with cell type–resolved mouse brain proteomics, and were further extended to represent the tripartite synapse. Enzyme-constrained resource allocation models (RAMs) were reconstructed for the astrocyte and neuron models using the GECKO 3.0 pipeline and integrated into a tripartite synapse community model via a modified SteadyCom framework. In the stoichiometry-only model, minimum lactate exchange remained at zero under ATP-maintenance and biomass objectives, indicating no coupling between astrocyte lactate secretion and neuronal uptake arising from the metabolic network itself. Imposing enzyme constraints made lactate secretion obligatory, with corresponding neuronal lactate uptake. Across 100 turnover-number bootstrap parameterizations, lactate coupling became obligatory under both objectives, with median onsets at 0.770 of maximal ATP-maintenance demand and 0.879 of maximal growth. This indicates that protein limitation can drive astrocyte-neuron lactate coupling consistent with ANLS, independent of modeling assumptions. Incorporation of reactive oxygen species (ROS) and lipid peroxidation pathways demonstrated that the brain-type fatty acid binding protein (FABP7) knockout redistributed flux away from astrocytic detoxification, increased neuronal ROS burden, and produced cell-type-specific changes in lipid-associated reporter metabolites, with prostaglandin J2 as the highest-ranked reporter metabolite. Model predictions, including lactate shuttling and lipid metabolic shifts in FABP7 knockout, align with published experimental studies, supporting the biological relevance of this framework. These findings indicate that ANLS emerges, at least in part, from enzyme limitation in the tripartite synapse model and that FABP7-mediated lipid trafficking is essential for lipid–redox homeostasis at the tripartite synapse.

**Author Summary:** The brain relies on metabolic cooperation between astrocytes and neurons. Astrocytes convert glucose into lactate, which neurons use for energy. This process is known as the astrocyte– neuron lactate shuttle (ANLS). It is still unclear whether this shuttle is just one possible metabolic pathway or is required due to resource limitations. To explore this, we created two detailed mathematical models of the tripartite synapse, where an astrocyte process extends to meet the synapse. The first model consisted only of the stoichiometry of the metabolic network, whereas the second reflected each cell’s limited ability to make proteins. When these protein limits were imposed, lactate secretion by astrocytes and uptake by the neurons became mandatory, providing a systems-level mathematics-driven demonstration of ANLS. We also examined the role of Fatty Acid Binding Protein 7 (FABP7), a protein primarily expressed in astrocytes that shuttles fatty acids within astrocyte cells. Modeling FABP7 knockout reduced astrocytes ROS detoxification capacity to 3.7% of wild type, increased oxidative stress in neurons, and produced cell-type-specific changes in lipid-associated reporter metabolites. These results show how energy supply, lipid trafficking, and oxidative stress control are linked across cell types and provide a way to identify metabolic vulnerabilities in neurological diseases.

## Introduction

The mammalian brain depends on tightly coordinated metabolic interactions between neurons and astrocytes to sustain synaptic transmission (communication between neurons), restore membrane polarization (returning cells to their resting electrical state), and buffer oxidative stress (protect cells from damage caused by reactive oxygen species) [1–4]. Neurons exhibit high oxidative demand but possess limited metabolic flexibility, as they store minimal glycogen and have a restricted ability to utilize alternative energy substrates under metabolic stress [1, 5]. Astrocytes compensate by providing metabolic support, recycling neurotransmitters, and detoxifying peroxidized lipids produced by neuronal oxidative stress, as shown in Fig 1A [3–6]. Disruption of the neuron–astrocyte metabolic interactions impair neuronal homeostasis and is linked to neurological disorders, including epilepsy, Alzheimer’s disease, traumatic brain injury, and cerebral ischemia [7–10].

**Fig 1.**
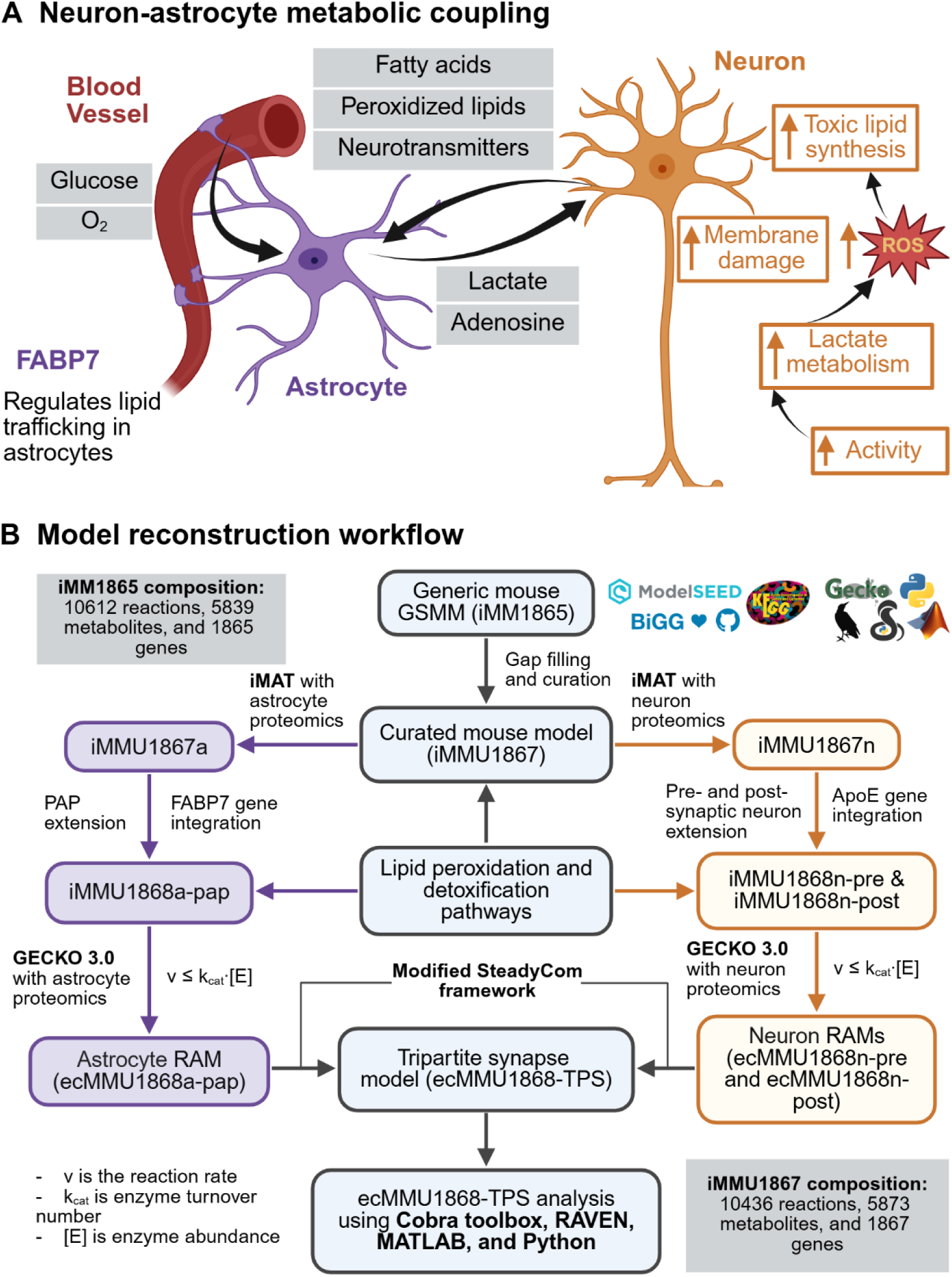
Neuron–astrocyte metabolic coupling and model reconstruction workflow. (A) Schematic of neuron– astrocyte metabolic coupling. Astrocytes take up glucose and O₂ from the blood vessel and supply lactate to neurons via the astrocyte–neuron lactate shuttle (ANLS). Neurons rely on astrocyte-derived lactate to meet energy demands, while astrocytes regulate redox homeostasis by buffering reactive oxygen species (ROS) and detoxifying lipid peroxidation products. Peroxidized lipids generated in neurons are transferred to astrocytes, where fatty acid-binding protein 7 (FABP7) facilitates intracellular trafficking and supports lipid detoxification and metabolic processing in the astrocyte cytosol. (B) Model reconstruction workflow. iMM1865 was gap-filled and curated to obtain iMMU1867. Cell-type-specific astrocyte (iMMU1867a) and neuron (iMMU1867n) models were generated using iMAT with neuron- and astrocyte-specific proteomics data. iMMU1867a was extended to include the PAP compartment (iMMU1868a-pap) and the FABP7 gene. Two copies of iMMU1867n were created, one with a pre-synaptic compartment (iMMU1868n-pre) and the other with a post-synaptic compartment (iMMU1868n-post) and the ApoE gene. Lipid peroxidation and detoxification pathways were added to all three models. Enzyme-constrained resource allocation models (RAMs) were reconstructed with GECKO 3.0, imposing *vj* ≤ *kcat*,*j* ⋅ [*Ej*], and coupled into a tripartite synapse community model via a modified SteadyCom framework. The community model was used for wild-type vs. FABP7-knockout comparisons and ANLS analysis. Databases: BiGG, ModelSEED, KEGG. Tools: COBRApy, RAVEN, GECKO, MATLAB, Python. Created in BioRender. Schroeder, W. (2026) https://BioRender.com/0dzvzht

The astrocyte–neuron lactate shuttle (ANLS) hypothesis provides a central framework for understanding metabolic coupling between astrocytes and neurons. According to this hypothesis, astrocytes respond to neuronal activity by metabolizing glucose to lactate and exporting it to neurons, where lactate serves as a preferred oxidative substrate for mitochondrial ATP production (Fig 1A) [6, 11, 12]. This hypothesis is supported by substantial experimental evidence, including studies demonstrating that neurons preferentially utilize lactate produced by astrocytes and depend on it for survival under stress [5, 6, 11, 12]. However, it remains unclear whether ANLS is universally required under physiological conditions or if it emerges only under specific metabolic constraints [1]. One unresolved possibility is that ANLS emerges when mitochondrial protein capacity becomes limiting, forcing astrocytes toward overflow metabolism. This scenario parallels overflow metabolism observed in microorganisms, where enzyme capacity limitations force cells to use less energy-efficient but more proteome-efficient pathways with higher throughput, even when more efficient routes are available [13, 14]. Enzyme-constrained metabolic models incorporate catalytic capacity and protein allocation, providing a framework for testing this hypothesis [14, 15].

Lactate shuttling is also functionally linked to redox and lipid metabolism. Neuronal oxidative metabolism generates reactive oxygen species (ROS), which promote lipid peroxidation and the accumulation of toxic lipid intermediates [5, 12, 16]. Astrocytes mitigate this oxidative burden by importing peroxidized lipids from neurons and process them through β-oxidation, lipid droplet formation, and detoxification pathways [5, 12, 16]. This interplay indicates that ANLS operates as part of a broader cross-cellular homeostatic system that coordinates energy supply with oxidative stress regulation.

The brain-type fatty acid binding protein (FABP7) is an astrocyte-enriched lipid chaperone that facilitates intracellular fatty acid trafficking and protects against ROS toxicity by regulating lipid handling and lipid droplet formation (Fig 2) [17–20]. As FABP7 operates at the intersection of lipid transport and detoxification within astrocytes, it serves as a biologically relevant model for investigating how disruptions in lipid handling reshape neuron–astrocyte coupling. Dysregulation of FABP7 has been implicated in various neurological disorders, including epilepsy, Alzheimer’s disease, and traumatic brain injury, highlighting its role in neuronal health and disease states [18, 20]. In addition to the ANLS hypothesis, this study examines FABP7 as a lipid-trafficking regulator and evaluates its contribution to redox homeostasis at the tripartite synapse, linking lipid metabolism to neuron–astrocyte coupling and the ANLS.

**Fig 2.**
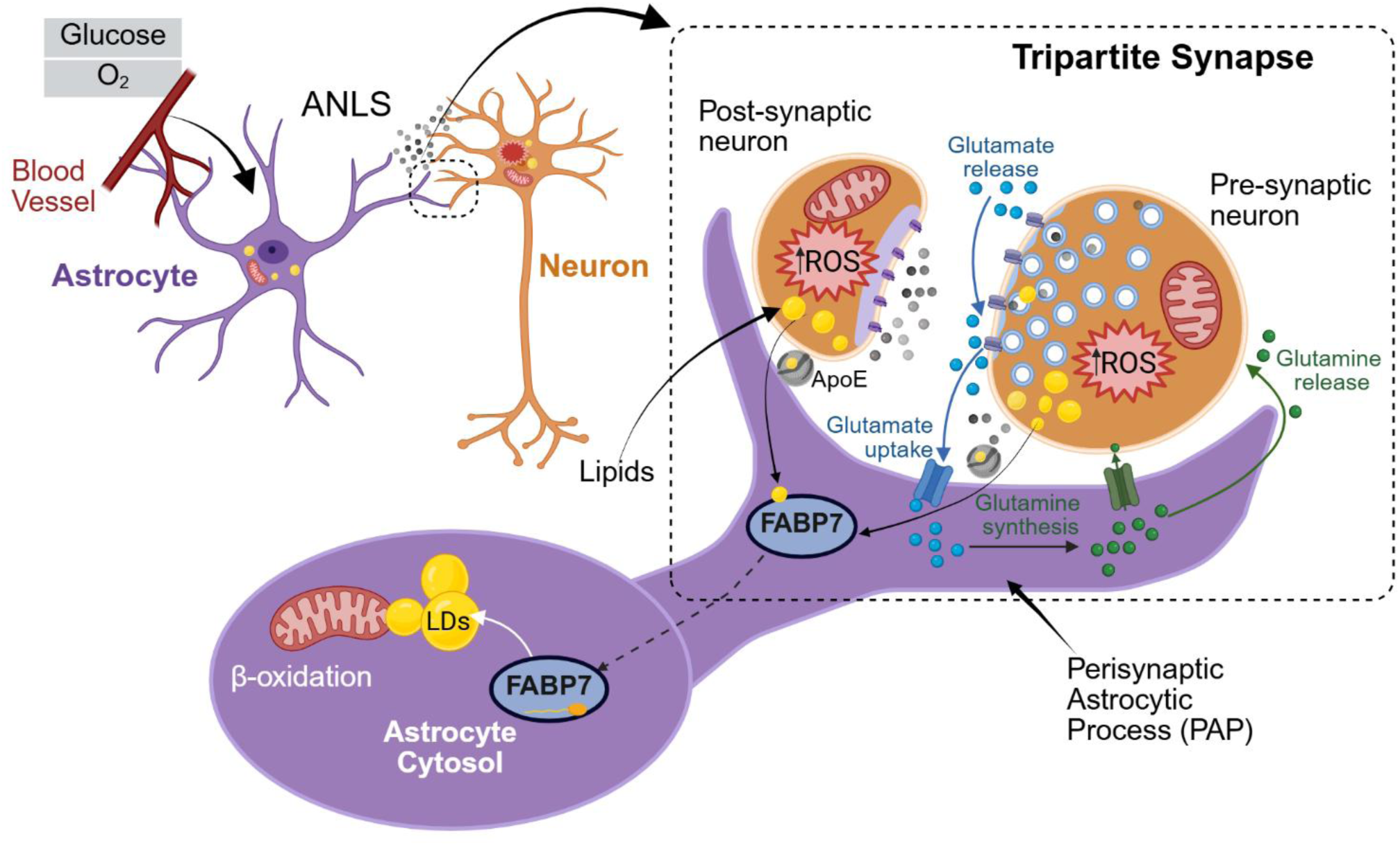
Mechanistic representation of the tripartite synapse model. Astrocytes take up glucose and O₂ from the blood vessel and supply lactate to neurons via the astrocyte–neuron lactate shuttle (ANLS). The model also represents glutamate release from the presynaptic neuron, glutamate uptake into the PAP, and conversion of glutamate to glutamine, which is transported back to the presynaptic neuron to support continued glutamate synthesis. Neuronal activity generates reactive oxygen species (ROS), leading to lipid peroxidation and the export of peroxidized lipids to the synaptic cleft via apolipoprotein E (ApoE)-mediated transport, where they are taken up into the PAP. Fatty acid-binding protein 7 (FABP7) then mediates transfer of these lipids from the PAP into the astrocyte cytosol, where they are incorporated into lipid droplets (LDs) and processed through detoxification and metabolic pathways. Stored lipids can subsequently be utilized by astrocyte mitochondria via β-oxidation to generate metabolic substrates, including ketone bodies. Created in BioRender. Schroeder, W. (2026) https://BioRender.com/et89oqf

Genome-scale metabolic models (GSMMs) provide a quantitative framework for analyzing complex, large, and multiscale metabolic interactions [14, 21, 22]. Previous genome-scale reconstructions of human brain metabolism demonstrated the feasibility of multicellular metabolic modeling in the central nervous system [23, 24]. However, these models did not capture the complexity of the tripartite synapse, investigate the mechanistic basis of ANLS, or consider protein limitations in key phenomena like the ANLS and redox balancing. This study addresses two major gaps: the need for a high-quality mouse metabolic reconstruction and the lack of an enzyme-constrained tripartite synapse community model for investigating ANLS and neuron–astrocyte metabolic coupling. Enzyme-constrained extensions of GSMMs, known as resource allocation models (RAMs), incorporate catalytic capacity and protein abundance limitations to provide more physiologically accurate predictions than stoichiometric models alone [14, 15]. In this study, the mouse metabolic model iMM1865 [25] was curated to address deficiencies in fatty acid and lipid pathways, stoichiometric consistency, and gene–reaction annotations. This new curated stoichiometric model is named iMMU1867. Astrocyte-specific (iMMU1867a) and neuron-specific (iMMU1867n) reconstructions were derived from the curated iMMU1867 model using iMAT and cell-type–resolved mouse brain proteomics data [26, 27]. The models were expanded to represent a tripartite synapse architecture, including a perisynaptic astrocytic process (PAP) and pre- and post-synaptic neuronal compartments to create a stoichiometric-only (iMMU1868-TPS) representation of the tripartite synapse linked via a modified SteadyCom framework [28]. The resulting stoichiometric models (astrocyte, iMMU1868a-pap; pre-synaptic neuron, iMMU1868n-pre; and post-synaptic neuron, iMMU1868n-post) were used as inputs to the GECKO 3.0 pipeline [15] to reconstruct enzyme-constrained RAMs of astrocytes (ecMMU1868a-pap) and neurons (ecMMU1868n-pre and ecMMU1868n-post) as shown in Fig 1B. Simulation of the enzyme-constrained tripartite synapse community model (ecMMU1868-TPS) demonstrated that enzyme limitation alone is sufficient to generate obligate lactate overflow consistent with ANLS, and that FABP7-mediated lipid trafficking is the dominant route maintaining lipid–redox homeostasis at the tripartite synapse. **Results**

The reconstruction proceeded through five main stages, with S1 Table summarizing the model output, data inputs, key assumptions, parameter values, and validation checks associated with each stage.

### Curation of iMM1865 resolves stoichiometric inconsistencies and improves model quality

Manual curation of the iMM1865 mouse metabolic model produced iMMU1867, which serves as the foundation for subsequent cell-type-specific and tripartite synapse modeling (see Fig 1B and Methods for detailed model curation and reconstruction). The curated network comprises 10,436 reactions, 5,873 metabolites, and 1,867 genes, representing a net change of −176 reactions, +36 metabolites, and +2 genes relative to iMM1865. These changes eliminated redundant lipid isoforms (R-groups) and improved the representation of brain lipid metabolism.

Fatty acid biosynthesis was expanded through the introduction of acyl-carrier protein (ACP) and acyl-CoA intermediates spanning C4-C18 chain elongation (KEGG map00061) [29], thereby restoring missing steps in the iMM1865 fatty acid elongation pathway. This reconstruction connects fatty acid synthesis intermediates to pooled lipid reactions, allowing newly synthesized fatty acids to be incorporated into membrane lipid synthesis. The addition of 3-hydroxyacyl-thioester dehydratase 2 (Entrez Gene ID: 109729085) and oleoyl-ACP (Entrez Gene ID: 99035) hydrolase genes enabled the completion of previously absent steps.

The curation process also addressed unrealistic exchange reactions that led to non-physiological growth. Numerous exchange reactions permitting unbounded uptake of intracellular metabolites were constrained to physiologically realistic bounds. That is, all 1,563 exchange reactions in iMM1865 used the SBML default lower bound of −1000 mmol·gDW⁻^1^·h⁻^1^, permitting unlimited uptake of every exchange metabolite. FBA under these bounds yields a biomass flux of 798.81 h⁻^1^ (Table 1), corresponding to a doubling time of approximately 3 seconds. To address this issue, a defined minimal medium was formulated from basic biomass-supporting nutrients, including glucose, oxygen, inorganic ions, phosphate, ammonium, and essential amino acids (S2 Table). The medium composition was selected to support mammalian cell growth, while uptake bounds were parameterized to approximate the astrocyte biomass flux range reported in a previous modeling study, including an experimental biomass flux of 0.32 h⁻^1^ and a model-predicted value of 0.33 h⁻^1^ [30]. Under these revised conditions, flux balance analysis (FBA) with biomass production as the objective yielded a growth rate of 0.327 h⁻^1^ in iMMU1867. Uptake bounds were calibrated against the reference values described above, and the medium was adjusted to reproduce the target growth rate. This rate exceeds measured brain macromolecular turnover and serves as a calibration reference rather than a physiological turnover rate. Because mature astrocytes and neurons are largely non-proliferative [30–32], biomass production flux is interpreted as macromolecular turnover and cell homeostasis rather than cell division. Non-dividing brain cells continuously replace their constituents over days to weeks [33–35]. This replacement requires continuous synthesis of the same macromolecular components that constitute the cell; thus, a composition-derived biomass reaction provides a reasonable approximation of the aggregate biosynthetic demand associated with cellular maintenance and turnover. Accordingly, ATP-maintenance (ATPM) was used as the primary objective for most subsequent analyses.

**Table 1.** Carbon balance comparison between iMM1865 and iMMU1867 under the defined medium, showing carbon uptake, carbon secretion, biomass incorporation, and residual imbalance. Full carbon analysis details are provided in S5 Table.

| Metric | iMM1865 | iMMU1867 (curated) |
| --- | --- | --- |
| Biomass ($\text{h}^{-1}$) | 798.811 | 0.327 |
| Carbon-bearing uptake exchanges | 30 | 10 |
| Carbon-bearing secretion exchanges | 40 | 5 |
| Total C input ($\text{C}\cdot\text{mmol}\cdot\text{gDW}^{-1}\cdot\text{h}^{-1}$) | 158,550.8 | 34.1 |
| Total C output ($\text{C}\cdot\text{mmol}\cdot\text{gDW}^{-1}\cdot\text{h}^{-1}$) | 239,312.2 | 15.7 |
| C to biomass ($\text{C}\cdot\text{mmol}\cdot\text{gDW}^{-1}\cdot\text{h}^{-1}$) | 30,256.5 | 17.0 |
| Carbon residual (IN – OUT – biomass) | –111,018.9 (33.7%) | +1.45 (4.2%) |
| Dominant uptake | distributed across 30 uptake exchanges | glucose (83.5%) |
| Dominant secretion | distributed across 40 secretion exchanges | lactate (73.9%) + $\text{CO}_2$ (24.4%) |

The MEMOTE report of iMM1865 identified 290 mass- and charge-imbalanced reactions, which were resolved by updating molecular formulas, metabolite charges, and reaction stoichiometry, substantially improving mass and charge balance. The implementation of a weighted-average fatty-acid pool reaction (modeled as FATTOTAL), calculated from experimental mouse brain lipidomics data [36], addressed the distribution of R-group fatty-acid composition in phospholipid reactions while maintaining stoichiometric consistency. Consequently, iMMU1867 passes MEMOTE’s stoichiometric consistency check, with mass and charge balance reaching 99.95% and 100%, respectively (S3 Table). Four reactions remain flagged as mass-imbalanced due to unresolved structural representations of biomass components, as detailed in S3 Table.

Gene–reaction associations and metabolite annotations were standardized, and Systems Biology Ontology (SBO) terms were comprehensively assigned. SBO annotation coverage increased from 0% to 78.8%. Together with the mass, charge, stoichiometric, and reaction boundary corrections described above, these improvements raised the overall MEMOTE [37] score from 46% to 83% (Fig 3A). A complete comparison between iMM1865 and iMMU1867 is provided in S3 Table, and the corresponding MEMOTE reports for both models are available in the GitHub repository (https://github.com/SchroederLabWSU/ecMMU1868-TPS).

**Fig 3.**
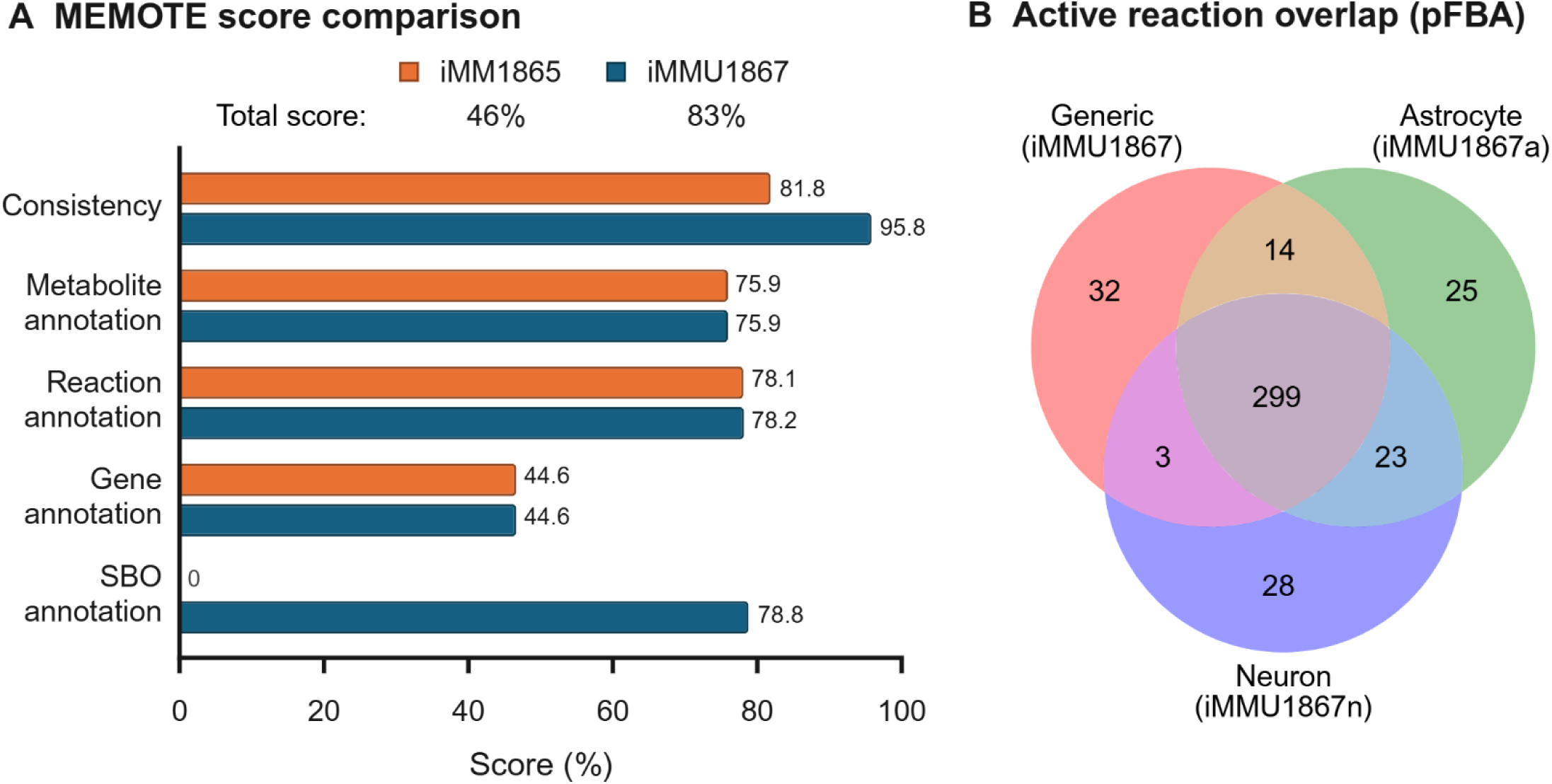
Model quality improvement and active reaction overlap following curation and cell-specific reconstruction. (A) MEMOTE score comparison between iMM1865 and iMMU1867 across consistency, annotation, and Systems Biology Ontology (SBO) categories. Curation improved model quality, increasing the final MEMOTE score from 46% in iMM1865 to 83% in iMMU1867. The largest improvement occurred in SBO annotation, which increased from 0% to 78.8% because the starting model lacked SBO terms. The consistency score also increased from 81.8% to 95.8%, reflecting improvements in mass and charge balancing. (B) Active reaction overlap across the generic model (iMMU1867) and cell-specific reconstructions (iMMU1867a and iMMU1867n), determined using pFBA. A total of 299 reactions are shared among all three models. The astrocyte model contains 25 unique reactions, the neuron model contains 28 unique reactions, and 32 reactions are unique to iMMU1867, highlighting both cell-type-specific metabolic specialization and core metabolic functions. See S6 Table for a complete list of active reactions and overlap reactions.

### Phenotypic validation

Four phenotypic tests confirmed physiologically consistent behavior of iMMU1867 (Fig 4 and S4 Table). Substrate utilization was evaluated by supplying glucose, lactate, glutamine, palmitate, or β-hydroxybutyrate (β-OHB) individually. Growth saturated for all five substrates at 22 mmol C gDW^⁻1^·h^⁻1^. Below this saturation point, growth rates varied among substrates and generally corresponded to their degree of reduction [38], with palmitate supporting the highest growth and glutamine the lowest. β-OHB supported slightly lower growth than glucose or lactate despite its higher degree of reduction, because its carbon enters central metabolism solely as acetyl-CoA [39]. (Fig 4A).

**Fig 4.**
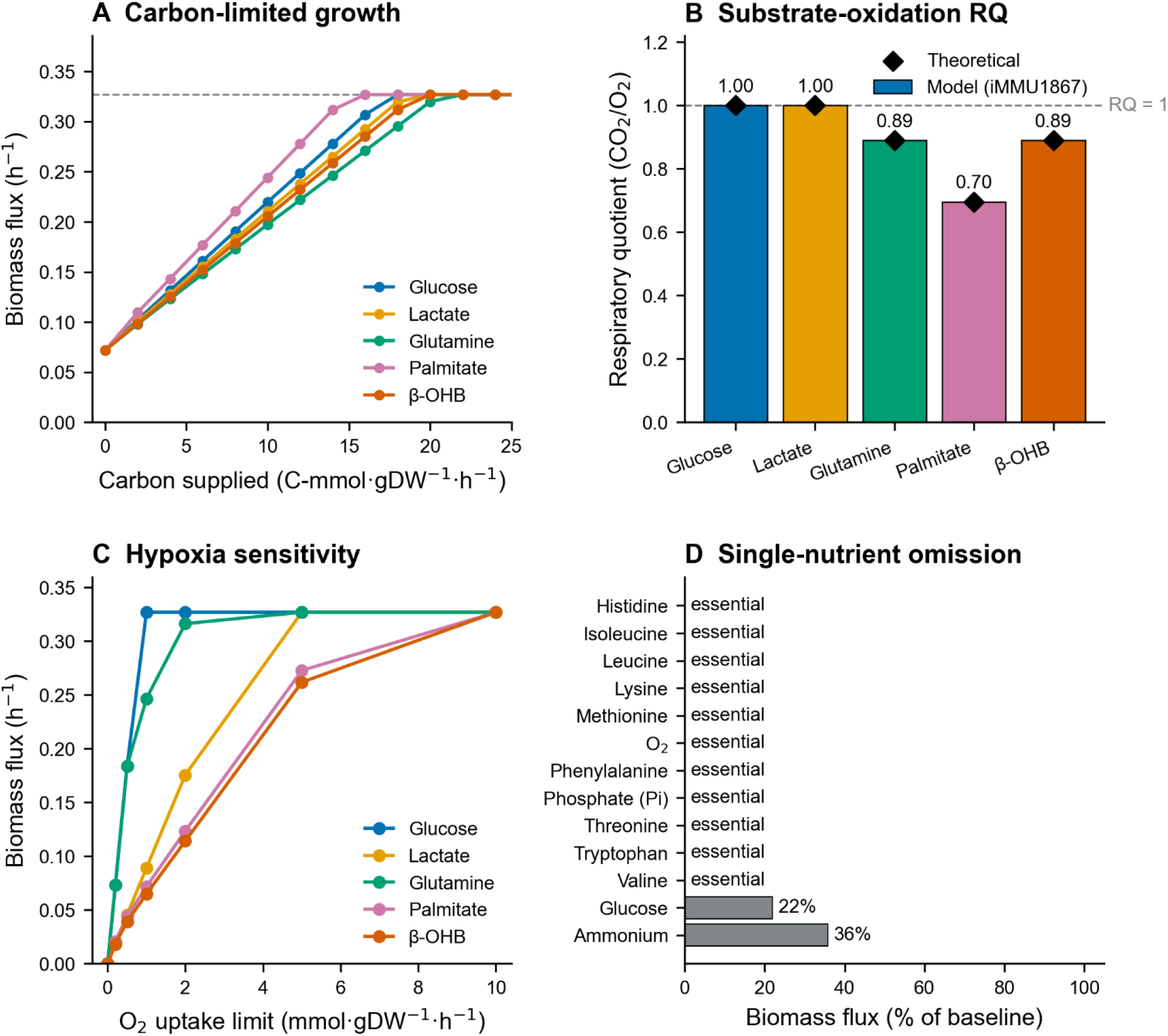
Phenotypic validation of iMMU1867. (A) Maximum biomass flux as a function of carbon supplied, with each substrate supplied alone in the defined minimal medium. All five substrates converged to the same maximum biomass flux of 0.327 h⁻¹ but saturated at different carbon supplies: palmitate at 16, glucose at 18, β-OHB and lactate at 20, and glutamine at 22 C-mmol·gDW^⁻1^·h^⁻1^. Below saturation, the substrates produced distinct growth rates, with palmitate supporting the highest biomass flux, followed by glucose, lactate, β-OHB, and glutamine.(B) Substrate-oxidation respiratory quotient (RQ = CO₂ secretion / O₂ uptake) under carbon-isolated conditions. Bars show iMMU1867-predicted RQ values, while black diamonds indicate theoretical complete-oxidation RQ values. The model reproduced the expected RQ values for glucose, glutamine, lactate, β-hydroxybutyrate (β-OHB), and palmitate. (C) Hypoxia sensitivity sweep across O₂ uptake limits of 10, 5, 2, 1, 0.5, 0.2, and 0 mmol·gDW^⁻1^·h^⁻1^. Glucose maintained maximal growth down to an O₂ uptake of 1 mmol·gDW^⁻1^·h^⁻1^ and was the most hypoxia-tolerant substrate, while glutamine retained near-maximal growth down to 2 mmol·gDW^⁻1^·h^⁻1^. Lactate maintained maximal growth down to 5 mmol·gDW^⁻1^·h^⁻1^ but declined at lower oxygen availability, whereas growth on palmitate and β-hydroxybutyrate declined once O₂ uptake fell below 10 mmol·gDW^⁻1^·h^⁻1^. All substrates showed no growth under anoxic conditions. (D) Single-nutrient omission identifies 11 essential exchanges (O₂, phosphate, and the nine essential amino acids). Glucose omission reduces biomass to 22% of baseline and ammonium to 36%, identifying both as growth-limiting but non-essential.

Substrate oxidation was further evaluated using respiratory quotient (RQ = CO_2_ secretion/O_2_ uptake), computed by maximizing ATP-maintenance (ATPM) under carbon-isolation conditions. iMMU1867 recapitulated the theoretical complete-oxidation RQ for glucose (1.00), lactate (1.00), glutamine (0.89), β-hydroxybutyrate (0.89), and palmitate (0.70), consistent with classical indirect calorimetry and principles of substrate oxidation [40, 41] (Fig 4B). The lower RQ values for β-hydroxybutyrate and palmitate reflect their greater oxygen requirement during oxidation, consistent with established fatty acid and ketone body metabolism [39, 42]. By contrast, the same substrate-oxidation test failed in the iMM1865, which reached the ATPM upper bound of 1000 mmol·gDW⁻^1^·h⁻^1^ with O_2_ uptake of 302.63 mmol·gDW⁻^1^·h⁻^1^ but no CO_2_ secretion across all five substrates (S4 Table). These results further confirm that curation removed non-physiological energy-generating cycles and unconstrained exchanges.

Oxygen limitation revealed substrate-specific differences in hypoxia tolerance (Fig 4C). Glucose maintained maximal growth down to an O_2_ uptake of 1 mmol·gDW⁻^1^·h⁻^1^ and was the most hypoxia-tolerant substrate, while glutamine retained near-maximal growth down to 2 mmol·gDW⁻^1^·h⁻^1^. Lactate maintained maximal growth down to 5 mmol·gDW⁻^1^·h⁻^1^ but declined at lower oxygen availability, whereas growth on palmitate and β-hydroxybutyrate declined once O_2_ uptake fell below 10 mmol·gDW⁻^1^·h⁻^1^. Glucose showed the highest hypoxia tolerance among the tested substrates because glycolysis produces ATP independently of oxygen. Glutamine ranked second, supporting mitochondrial substrate-level phosphorylation via succinyl-CoA synthetase and entering the tricarboxylic acid cycle through α-ketoglutarate [43, 44]. Under limited oxygen conditions, succinyl-CoA synthetase facilitated substantial ATP production from glutamine, minimal production from lactate, and none from glucose. Although lactate sustained greater oxidative phosphorylation than glutamine, it supported lower cellular growth, suggesting that glutamine’s hypoxia tolerance is primarily attributable to substrate-level phosphorylation. Palmitate and β-OHB exhibited lower hypoxia tolerance because their carbon enters central metabolism mainly as acetyl-CoA, which requires mitochondrial oxidative metabolism. Additionally, fatty acid oxidation incurs a higher oxygen cost per ATP generated compared to glucose [45]. No substrate-supported growth occurred under anoxic conditions, which aligns with the established oxygen dependence of brain metabolism [10, 46].

In Fig 4D, a single-nutrient omission test on the defined medium (S2 Table) identified 11 essential exchanges, including O_2_, phosphate, and the nine essential amino acids (Phe, Met, Lys, Leu, His, Ile, Val, Thr, Trp), consistent with known mammalian nutrient requirements [47, 48]. Glucose omission reduced biomass to 22% of baseline, while ammonium reduced it to 36%, indicating that both are strongly growth-limiting but not essential. Both are non-essential due to the re-direction of essential amino acids into pathways for the biosynthesis of macromolecules and biomass precursors.

Collectively, these results indicate that iMMU1867 reproduces physiologically-consistent metabolic behavior and provides a robust foundation for subsequent mechanistic analyses of neuron–astrocyte metabolic coupling.

### Carbon balance analysis quantifies the impact of model curation

Following phenotypic validation, network-level carbon balance analysis showed that curation substantially reduced non-physiological carbon flux (Table 1 and S5 Table). Exchange reactions in the SBML model were identified using regular-expression searches, confirming that the base model permitted unbounded exchange fluxes and, consequently, unrealistic nutrient uptake. Carbon balance analysis was subsequently employed to compare total carbon uptake with total carbon secretion, thereby providing a network-level assessment of non-physiological carbon exchange while maintaining elemental consistency. In the iMM1865 mouse model, FBA with biomass objective yielded an unrealistically high growth rate of 798.81 h⁻^1^, driven by excessive, unbounded exchange uptake fluxes. Carbon flowed through 30 uptake exchange reactions contributed 158,551 C-mmol·gDW⁻^1^·h⁻^1^, while 40 secretion exchanges and biomass synthesis accounted for 239,312 C-mmol·gDW⁻^1^·h⁻^1^ (Table 1). This imbalance resulted in a gain of 111,018 C-mmol·gDW⁻^1^·h⁻^1^ (+33.7% more than the uptake of carbon), indicating a substantial violation of the carbon balance. The dominant non-physiological carbon uptakes in iMM1865 were driven by exchange reactions including N-acylsphingosine (EX_crm_hs_e, 17,844 C-mmol·gDW⁻^1^·h⁻^1^), UDP-glucuronate (EX_udpglcur_e, 15,000 C-mmol·gDW⁻^1^·h⁻^1^), and nucleotide and nucleoside metabolites including ATP (EX_atp_e), xanthosine (EX_xtsn_e), and deoxythymidine-5’-phosphate (EX_dtmp_e), each contributing 10,000 C-mmol·gDW⁻^1^·h⁻^1^. These fluxes reflect unrealistic routes that allow carbon to enter and exit the network without physiologically realistic constraints.

Curation substantially reduced non-physiological carbon flux by restricting carbon uptake to the defined medium components listed in S2 Table. In iMMU1867, biomass production flux decreased to the reference value of 0.327 h⁻^1^ [30]. Within this defined medium, carbon uptake was constrained to 10 exchanges, dominated by glucose (83.5%) and the nine essential amino acids (16.5%). Under these input constraints, carbon secretion occurred through five main exchanges, primarily lactate (73.9%) and CO₂ (24.4%), with minor contributions from HCO₃⁻, 2-hydroxybutyrate, and succinate. Total carbon input was reduced to 34.1 C-mmol·gDW⁻^1^·h⁻^1^, with 15.7 C-mmol·gDW⁻^1^·h⁻^1^ secreted and 17.0 C-mmol·gDW⁻^1^·h⁻^1^ incorporated into biomass. The resulting residual of +1.45 C-mmol·gDW⁻^1^·h⁻^1^ indicates near-complete carbon closure, corresponding to a deviation of approximately 4.2% of total input (Table 1). This residual arises primarily from unresolved R-group representations in biomass lipid metabolites, including phosphatidylserine, phosphatidylethanolamine, phosphatidylglycerol, sphingomyelin, cardiolipin, and phosphatidylinositol, which were retained to preserve overall stoichiometric consistency.

### Cell-specific reconstruction and tripartite expansion establish a tripartite synapse modeling framework

Astrocyte-specific (iMMU1867a) and neuron-specific (iMMU1867n) reconstructions were derived from iMMU1867 by integrating cell-type-resolved mouse-brain proteomics [27] using iMAT [26], which incorporates expression data to identify the most likely active reaction set while preserving metabolic functionality. Both context-specific models retained the composition of iMMU1867 (10,436 reactions, 5,873 metabolites, 1,867 genes, and 9 compartments), differing primarily in reaction-bound constraints rather than reaction content. To assess the base metabolic differences between the cell-line specific models, active reaction sets were computed by parsimonious flux balance analysis (pFBA) [49]. The active reaction sets comprised 361 reactions in iMMU1867a and 353 in iMMU1867n, each supporting maximum biomass flux to that of iMMU1867 (0.327 h⁻^1^) (Table 2). Active-reaction overlap analysis with pFBA identified a conserved core of 299 shared reactions, with 25 reactions enriched in astrocytes (predominantly purine metabolism, peroxisomal transport, and cholesterol biosynthesis) and 28 reactions enriched in neurons (primarily phospholipid biosynthesis, nucleoside-diphosphate kinase metabolism, mitochondrial transport, and ROS detoxification) (Fig 3B). These differences recapitulate well-established cell-type metabolic specialization [4]. Complete active-reaction and overlap-reaction lists are provided in S6 Table.

**Table 2.** Stoichiometry-only genome-scale metabolic model statistics across reconstruction stages, including generic, cell-specific, and tripartite synapse reconstructions. Active reactions were identified from pFBA solutions.

| Model | Reactions | Metabolites | Genes | Compartments | Active reactions (pFBA) | Maximum growth rate (h<sup>-1</sup>) |
| --- | --- | --- | --- | --- | --- | --- |
| iMMU1867 (curated generic) | 10,436 | 5,873 | 1,867 | 9 | 348 | 0.327 |
| iMMU1867a (astrocyte) | 10,436 | 5,873 | 1,867 | 9 | 361 | 0.327 |
| iMMU1867n (neuron) | 10,436 | 5,873 | 1,867 | 9 | 353 | 0.327 |
| iMMU1868a-pap (astrocyte + PAP) | 10,465 | 5,895 | 1,868 | 11 | 363 | 0.327 |
| iMMU1868n-pre (pre-synaptic neuron) | 10,458 | 5,895 | 1,868 | 11 | 357 | 0.327 |
| iMMU1868n-post (post-synaptic neuron) | 10,455 | 5,886 | 1,868 | 11 | 354 | 0.327 |
| iMMU1868-TPS | 32,872 | 19,114 | 5604 (1868 unique genes) | 34 (11 per-cell + 1 shared synaptic cleft) | 816 | 0.105 |

To represent the tripartite synapse, iMMU1867a was expanded to include the perisynaptic astrocytic process (PAP), which ensheathes synapses (Fig 2). To further build a model of the tripartite synapse, the neuron model (iMMU1867n) was duplicated to model pre-synaptic and post-synaptic neurons. A synaptic cleft compartment (syn) was introduced to represent intercellular metabolite exchange at the synapse. Although both neuronal models share the same underlying reaction network, they were assigned distinct synaptic transport roles. The pre-synaptic neuron releases glutamate into the synaptic cleft and receives glutamine from the astrocytes for neurotransmitter recycling, whereas glutamine uptake is not permitted in the post-synaptic neuron. This reflects the different metabolic roles of pre- and post-synaptic neuronal compartments during synaptic transmission. This yielded three site-specific reconstructions: iMMU1868a-pap (astrocyte with PAP), iMMU1868n-pre (pre-synaptic neuron), and iMMU1868n-post (post-synaptic neuron). Each model contains 1,868 genes, reflecting the addition of apolipoprotein E (ApoE; Entrez Gene ID: 11816), which is required for lipid transport across the synaptic cleft [20]. Using pFBA, active reaction counts were 363 in iMMU1868a-pap, 357 in iMMU1868n-pre, and 354 in iMMU1868n-post, with all models maintaining a biomass flux value of 0.327 h⁻^1^ (Table 2).

Literature-supported intercellular transport reactions were incorporated to capture synaptic metabolism (S1 Data), including the astrocyte–neuron lactate shuttle, the glutamate–glutamine cycle, serine transport supporting N-methyl-D-aspartate (NMDA) receptor function, glutathione-mediated redox coupling, and ketone-body exchange via H⁺-coupled transport systems [50–54]. In parallel, a lipid peroxidation module was introduced to represent neuron-to-astrocyte transfer of damaged lipids (see S1 Data for complete reaction details). Reactive oxygen species (ROS) in neuronal compartments generate the peroxidized lipid species C04717 (13-L-hydroperoxylinoleic acid), which is represented in compartment-specific forms within the tripartite models. This species is exported from neurons to the synaptic cleft via ApoE-mediated transport (Entrez Gene ID: 11816), taken up into the PAP, and transferred into astrocyte cytosol by FABP7 (Entrez Gene ID: 12140) [12, 20, 55] (Fig 2). Within astrocytes, glutathione-dependent reduction and repair reactions detoxify and recycle the lipid species, completing a neuron–astrocyte lipid-handling pathway. This expansion establishes a mechanistically complete tripartite synapse framework suitable for enzyme-constrained and community-level analyses.

The stoichiometric models iMMU1868a-pap, iMMU1868n-pre, and iMMU1868n-post were linked using the SteadyCom framework [28] to create the stoichiometric reconstruction of the tripartite synapse, iMMU1868-TPS, as detailed in Methods.

### Enzyme constraints make ANLS obligatory across objectives and protein budgets

We reconstructed resource allocation models (RAMs) of astrocytes (ecMMU1868a-pap) and neurons (ecMMU1868n-pre and ecMMU1868n-post) using the GECKO 3.0 protocol [15] as shown in Fig 1B and detailed in Methods, using the stoichiometric models as the basis for reconstruction. The enzyme-constrained tripartite synapse community model (ecMMU1868-TPS) was constructed by combining the enzyme-constrained astrocyte, pre-synaptic neuron, and post-synaptic neuron models using the modified SteadyCom framework [28] described in the Methods. ecMMU1868-TPS consisted of 77,257 reactions, 24,661 metabolites, and 5,607 genes across 13 compartments per cell. The increased model size reflects the additional enzyme-associated components introduced during GECKO reconstruction.

To investigate if protein limitations drive the ANLS, we first compared the biomass-lactate production envelopes between the stoichiometric-only tripartite synapse community model (iMMU1868-TPS) and the enzyme-constrained tripartite synapse community model (ecMMU1868-TPS) (Fig 5A and 5B). This growth-based analysis was retained as the initial comparison, while ATP-maintenance was used as the primary physiological objective because mature astrocytes and neurons are largely non-proliferative [30–32].

**Fig 5.**
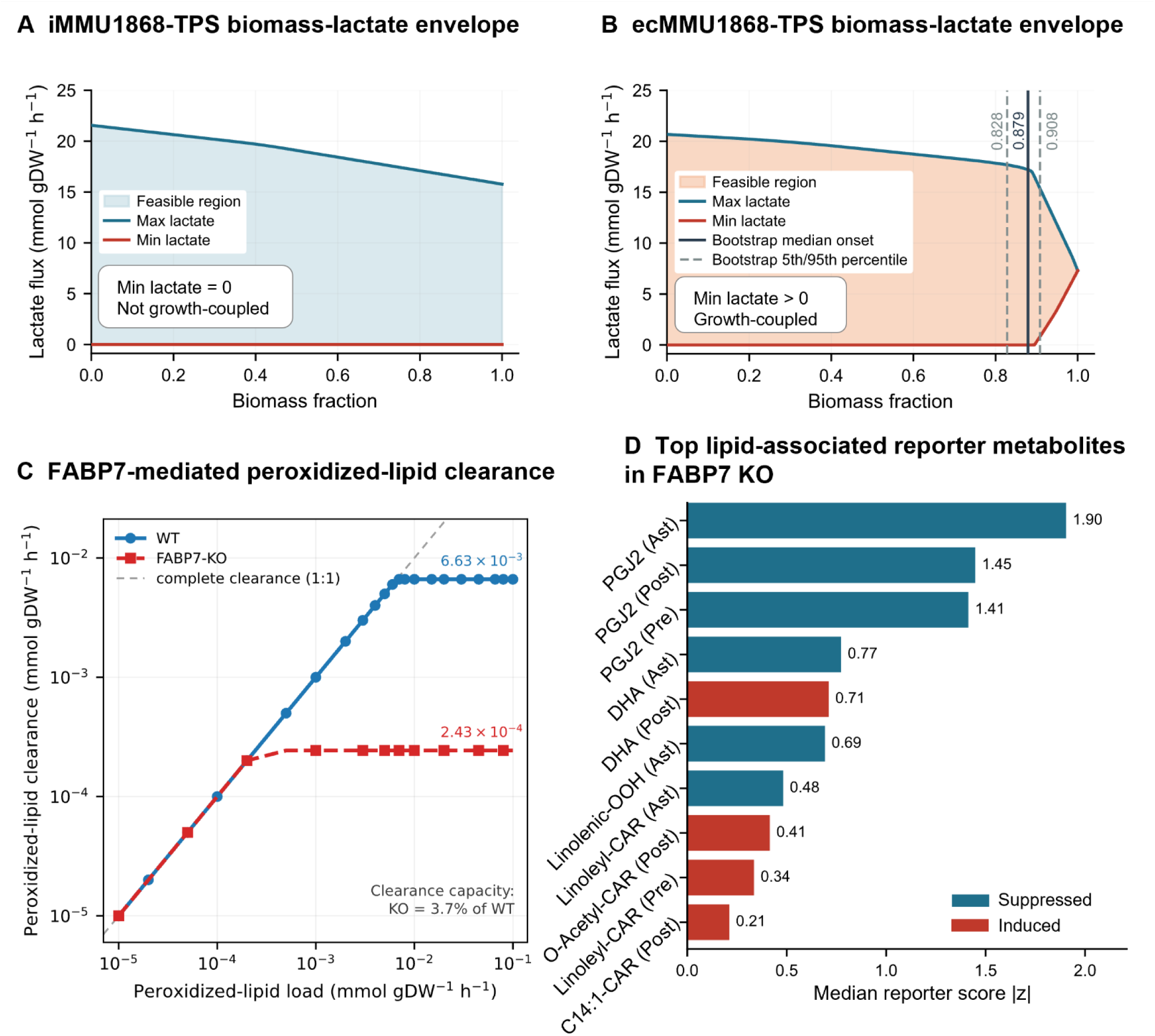
Lactate coupling and FABP7-dependent lipid/redox responses under the biomass objective in the tripartite synapse community model. (A) Biomass-lactate envelope of the stoichiometric TPS community model (iMMU1868-TPS). Minimum lactate secretion (orange) remains zero across the full biomass range, indicating that lactate export is not required for growth. Maximum lactate secretion (blue) decreases from approximately 21.6 to 15.8 mmol·gDW^⁻1^·h^⁻1^. The shaded region represents the feasible lactate secretion space. (B) Biomass-lactate envelope of the enzyme-constrained tripartite community model (ecMMU1868-TPS). Under the turnover numbers assigned by GECKO, lactate secretion becomes obligatory at 0.895 of maximal growth, indicating growth-coupled lactate overflow under enzyme-capacity constraints. The solid vertical line marks the median coupling onset across 100 *k*_cat_ bootstrap parameterizations (0.879), and the dashed lines indicate the 5th and 95th percentiles (0.828 and 0.908). Lactate secretion became obligatory in all 100 parameterizations. (C) WT and FABP7-knockout (KO) astrocytic clearance of neuron-derived peroxidized lipid (13-HPODE; C04717) under the biomass objective. Both genotypes completely cleared the delivered load up to approximately 2 × 10⁻⁴ mmol·gDW^⁻1^·h^⁻1^ (dashed line, 1:1). Above this load, KO clearance saturated at 2.43 × 10⁻⁴ mmol·gDW⁻¹·h⁻¹, whereas WT clearance continued to 6.63 × 10⁻³ mmol·gDW⁻¹·h⁻¹, with KO reaching 3.7% of WT clearance capacity at saturation.(D) Top lipid-associated reporter metabolites in FABP7-KO under biomass objective. Bars show the median reporter score |z| across 100 kcat bootstrap parameterizations for the ten highest-ranked lipid-associated reporter metabolites. Colors indicate suppressed or induced flux changes in the knockout. PGJ2, prostaglandin J2; DHA, docosahexaenoic acid; Linolenic-OOH, linolenic acid hydroperoxide; Linoleyl-CAR, linoleyl carnitine; C14:1-CAR, tetradecenoylcarnitine(5); Ast, astrocyte; Pre, presynaptic neuron; Post, postsynaptic neuron.

In iMMU1868-TPS, the maximum growth rate was 0.105 h⁻^1^. The minimum feasible lactate exchange (EX_lac L[u]) remained at zero across the entire biomass range, indicating that lactate secretion was optional and not coupled to community biomass production (Fig 5A). Introducing enzyme constraints reduced the maximum growth rate from 0.105 h⁻^1^ to 0.067 h⁻^1^ and reshaped the community lactate envelope. This reduction in growth is expected because enzymatic constraints limit the total proteome capacity available to support reaction fluxes, thereby reducing the feasible solution space relative to the stoichiometric model. Similar reductions in maximum biomass flux have been widely observed in enzyme-constrained and resource-allocation metabolic models following the incorporation of proteome limitations [13, 14, 56]. In ecMMU1868-TPS, minimum feasible lactate secretion remained zero across most of the biomass envelope and became obligatory at 0.895 of the maximal growth rate using the turnover numbers assigned by GECKO (Fig 5B and S7 Table). Across 100 turnover-number bootstrap parameterizations, lactate coupling became obligatory in all 100, with an onset between 0.828 and 0.908 of maximal growth (5th–95th percentile; median 0.879) (Fig 5B and S8 Table).

Lactate coupling was evaluated under the ATPM objective. Biomass production was fixed at zero in all three cell types, and the objective was defined as the summed ATPM flux in the astrocyte, presynaptic neuron, and postsynaptic neuron. No obligate lactate secretion was observed in the stoichiometric model under the ATPM objective (Fig 6A). In contrast, minimum feasible lactate secretion in ecMMU1868-TPS became obligatory above 0.83 of maximal ATP-maintenance demand using the GECKO-assigned turnover numbers (Fig 6B and S7 Table). Across the same 100 paired turnover-number bootstrap parameterizations, coupling became obligatory in all 100, with an onset between 0.660 and 0.840 of maximal demand (5th–95th percentile; median 0.770) (Fig 6B and S8 Table). The ATP-maintenance objective produced an earlier lactate-coupling onset than the biomass objective in every paired parameterization (S1 Fig), with median onsets of 0.770 and 0.879 of their respective maxima. At maximal maintenance demand, the two neuronal models together carried a median of 73% of the total ATPM flux, with the remainder in the astrocyte (S8 Table). This finding aligns with the energetics of neuronal signaling, where a major ATP cost is associated with restoring the transmembrane ion gradients dissipated by action potentials [57].

**Fig 6.**
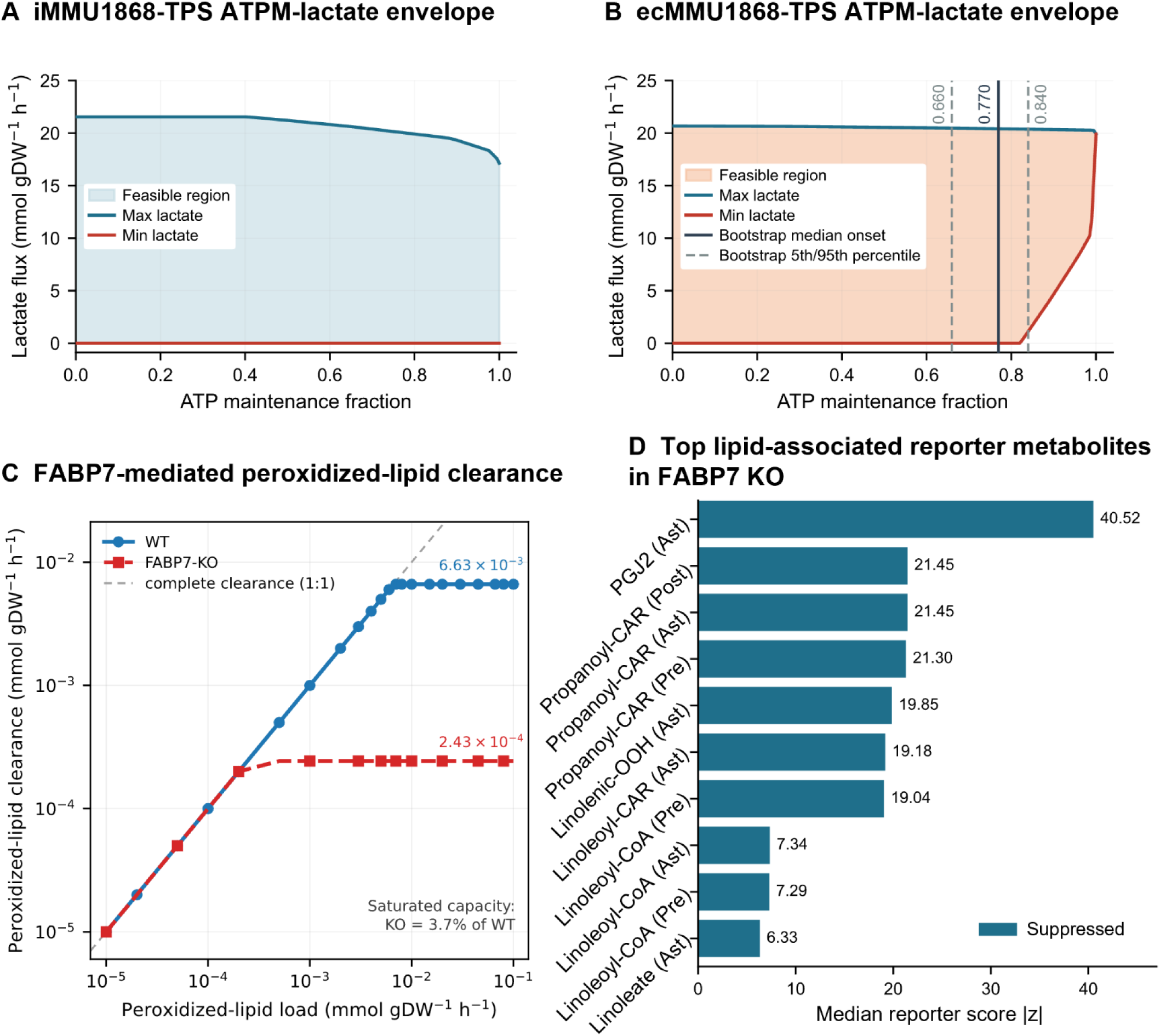
Lactate coupling and FABP7-dependent lipid/redox responses under the ATP-maintenance objective in the tripartite synapse community model. (A) ATP maintenance–lactate envelope of the stoichiometric community model (iMMU1868-TPS). Minimum lactate secretion remained zero across the full ATP-maintenance range, indicating that lactate export was not required to sustain maintenance demand. (B) ATP maintenance–lactate envelope of the enzyme-constrained community model (ecMMU1868-TPS). Under the turnover numbers assigned by GECKO, lactate secretion became obligatory at 0.825 of maximal ATP-maintenance demand. The solid vertical line marks a median coupling onset of 0.770 across the same 100 *k*_cat_ bootstrap parameterizations used in Fig 5B, and the dashed lines indicate the 5th and 95th percentiles (0.660 and 0.840). Lactate secretion became obligatory in all 100 parameterizations, with an earlier onset than under the biomass objective in every paired parameterization. (C) WT and FABP7-knockout (KO) astrocytic clearance of neuron-derived peroxidized lipid (13-HPODE) under the ATP-maintenance objective. WT clearance saturated at 6.63 × 10⁻³ mmol·gDW⁻¹ h⁻¹, whereas KO clearance saturated at 2.43 × 10⁻⁴ mmol·gDW⁻¹ h⁻¹, corresponding to 3.7% of WT capacity. These clearance capacities were unchanged from those obtained under the biomass objective (Fig 5C). (D) Lipid-associated reporter metabolites in FABP7-KO under the ATP-maintenance objective. Bars show the median reporter score |z| across 100 kcat bootstrap parameterizations for the ten highest-ranked lipid-associated reporter metabolites. All displayed metabolites were suppressed in the knockout. PGJ2, prostaglandin J2; Propanoyl-CAR, O-propanoylcarnitine; Linolenic-OOH, linolenic acid hydroperoxide; Linoleyl-CAR, linoleyl carnitine; Linoleoyl-CoA, linoleoyl coenzyme A; Ast, astrocyte; Pre, presynaptic neuron; Post, postsynaptic neuron.

We systematically varied the protein-pool bound across 25 values, ranging from 0.01× to 2.2× the baseline bound of 10 mmol·gDW^⁻1^·h^⁻1^ (0.1–22 mmol·gDW^⁻1^·h^⁻1^), and recomputed the lactate-coupling onset at each value. Obligatory lactate secretion was observed for bounds between 0.15× and 1.8×, with onset values ranging from 0.887 to 0.986 of maximal biomass flux (Fig 7A, regimes II-IV). Lactate coupling was absent below 0.15×, where the maximum feasible growth was insufficient to necessitate lactate overflow (Fig 7, regime I). Coupling was also lost above 1.8×, as all tested bounds from 1.9× to 2.2× exhibited the same behavior, with lactate secretion not being obligatory at any growth fraction (Fig 7A, regime V). Analysis of protein allocation clarified these regimes. Obligatory lactate secretion occurred only when the protein-pool constraint was binding, which persisted up to approximately 16.4 mmol·gDW^⁻1^·h^⁻1^ per cell. Beyond this threshold, the maximum growth rate could be achieved without utilizing the entire protein pool, resulting in the loss of obligatory lactate secretion at higher growth fractions (Fig 7, regime IV). The astrocyte reallocated resources from glycolysis toward glycerophospholipid metabolism, while the postsynaptic neuron shifted in the opposite direction. Both neuronal compartments increased allocation to folate-dependent metabolism, as well as serine and glycine metabolism (S9 Table). These results indicate that ANLS emerges under conditions of high metabolic demand due to protein-capacity limitation at the tripartite community level. Under these enzyme-constrained conditions, astrocytic lactate secretion is coupled to neuronal uptake through the shared extracellular pool (lac L[u]), consistent with the directional astrocyte-to-neuron lactate flux proposed by the ANLS framework. This lactate can support increased neuronal energetic demand, linking lactate overflow to neuronal energy support under high-growth, protein-limited conditions.

**Fig 7.**
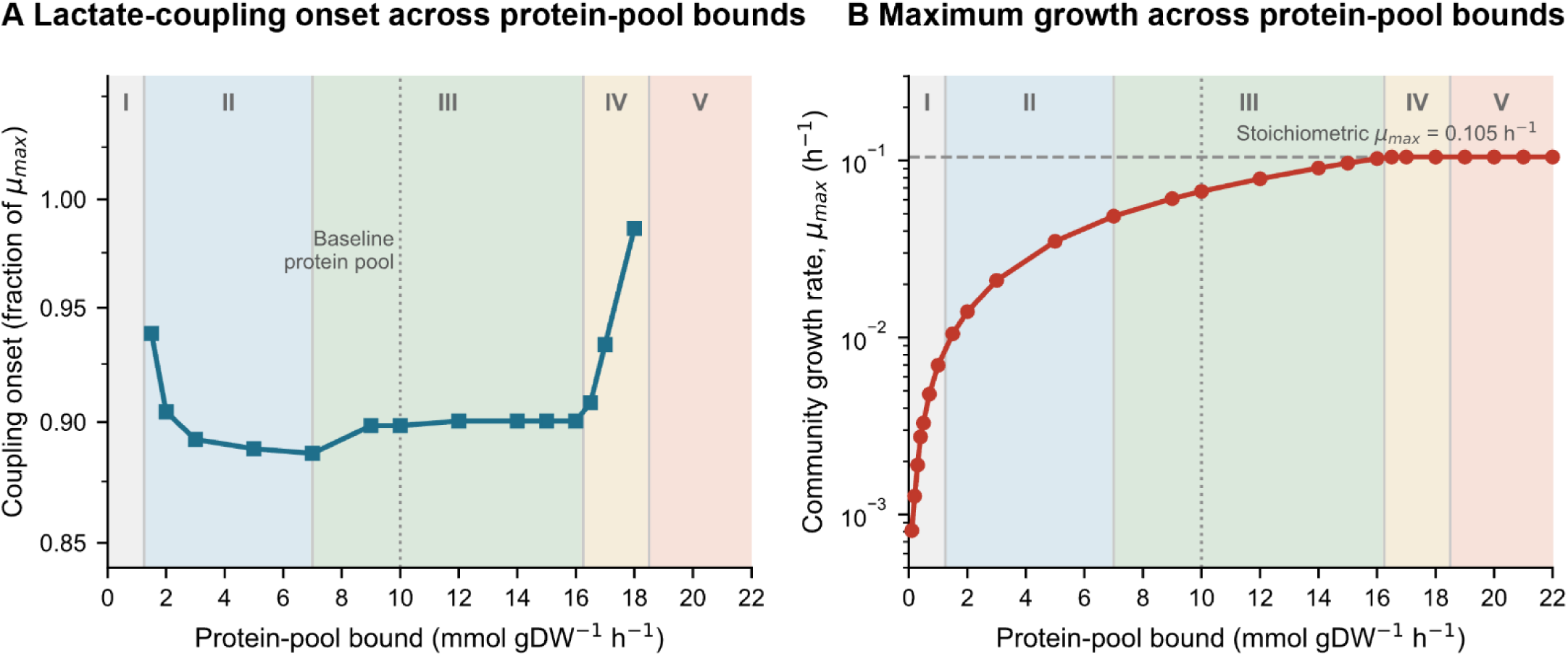
Lactate coupling and community growth across the enzyme protein-pool bound. The protein-pool bound was varied across 25 values from 0.1 to 22 mmol·gDW^⁻1^·h^⁻1^, corresponding to 0.01× to 2.2× the baseline bound of 10 mmol·gDW^⁻1^·h^⁻1^ (dotted vertical line). Shading indicates five regimes, I–V. (A) Lactate-coupling onset, defined as the lowest fraction of maximal growth at which lactate secretion becomes obligatory. Lactate coupling is absent in regime I, where maximum feasible growth is too low to require lactate overflow. In regime II, the onset decreases from 0.939 to 0.887 as the protein-pool bound increases, then remains near 0.90 throughout regime III. In regime IV, the onset rises sharply to 0.986, and in regime V lactate secretion is no longer obligatory at any growth fraction. (B) Maximum community growth rate across the same bounds, shown on a logarithmic axis. Growth increases with the protein-pool bound and reaches the stoichiometric ceiling of 0.105 h^⁻1^ (dashed horizontal line) near the III/IV boundary. The protein-pool constraint becomes nonbinding at approximately 16.4 mmol·gDW^⁻1^·h^⁻1^ per cell, after which the same maximum growth can be achieved without using the full protein pool and obligatory lactate secretion is progressively lost.

### FABP7-dependent lipid trafficking couples astrocytic detoxification to neuronal redox homeostasis

Wild-type (WT) and FABP7-knockout (KO) simulations in ecMMU1868-TPS were compared to determine the influence of astrocytic FABP7 on peroxidized lipid clearance and neuronal oxidative burden (Fig 5C and Fig 6C). To ensure FABP7 was not the sole route for peroxidized-lipid (13-HPODE; C04717) clearance, the lipid-trafficking module was expanded to include parallel FABP5- and FABP3-mediated transport pathways. FABP5 and FABP3 were parameterized based on their measured astrocytic abundance and DLKcat-predicted substrate-specific turnover numbers. These alternative pathways were present in both genotypes, and the knockout condition removed only FABP7-dependent reactions. Peroxidized-lipid clearance was quantified downstream of transport at the glutathione-dependent reduction step.

In this model, superoxide is converted to hydrogen peroxide, which is subsequently removed by catalase and glutathione-dependent reactions. Lipid peroxidation in neurons generates 13-HPODE, which astrocytes reduce using glutathione, thereby linking lipid detoxification to cellular redox metabolism. In WT simulations, peroxidized-lipid clearance increased with the delivered load until reaching saturation at 6.63 × 10⁻³ mmol·gDW⁻^1^·h⁻^1^. In the FABP7 knockout, clearance also increased with load but saturated at 2.43 × 10⁻⁴ mmol·gDW⁻^1^·h⁻^1^, representing 3.7% of WT capacity (Fig 5C). When the delivered load was below the residual FABP5/FABP3 capacity, the knockout cleared the load completely. The difference from WT became apparent only when the delivered load exceeded this residual capacity and increased further. Above this threshold, the knockout could no longer fully detoxify the delivered peroxidized lipid. Repeating the peroxidized-lipid clearance analysis under the ATPM objective resulted in no substantial change in clearance capacity. WT and FABP7-KO clearance saturated at 6.63 × 10⁻³ and 2.43 × 10⁻⁴ mmol·gDW⁻¹·h⁻¹, respectively, again leaving the knockout at 3.7% of WT capacity (Fig 6C).

Robustness analyses showed that, with community biomass fixed at its optimum, WT peroxidized-lipid clearance ranged from 1.00 × 10⁻³ to 6.63 × 10⁻³ mmol·gDW^⁻1^·h^⁻1^, whereas FABP7-KO clearance remained fixed at 2.43 × 10⁻⁴ mmol·gDW^⁻1^·h^⁻1^. Thus, the knockout maximum remained below the WT minimum across all alternative optimal solutions. Sensitivity analysis, in which FABP5 and FABP3 *k*_cat_ values were scaled from 0.1× to 10× their DLKcat predictions, increased knockout clearance only from 0.37% to 36.71% of WT capacity. Achieving full compensation would require approximately a 27-fold increase above the predicted turnover, which exceeds the ±1 log₁₀ range representing the reported DLKcat prediction error [58] (S10 Table and S2 Fig).

These results suggest that FABP7 serves as the primary modeled carrier for astrocytic peroxidized-lipid clearance, whereas FABP5 and FABP3 offer only limited residual compensation. Consequently, the loss of FABP7-mediated trafficking diminishes astrocytic detoxification capacity and elevates the modeled neuronal redox burden. A comprehensive load-dependent clearance analysis is presented in S10 Table.

### Reporter metabolite analysis identifies astrocytic prostaglandin J2 and linoleate-pathway suppression in the FABP7 knockout

Reporter-metabolite analysis [59], which identifies metabolites surrounded by coordinated changes in associated reaction fluxes, was performed to assess the impact of FABP7 knockout on the ecMMU1868-TPS community under conditions of forced peroxidized-lipid stress. ATP-maintenance served as the primary objective function, using the same 100 kcat bootstrap parameterizations as in the lactate coupling analysis. To improve biological interpretability, currency metabolites, GECKO enzyme pseudo-metabolites, biomass pools, and highly connected hub metabolites were excluded prior to scoring, and compartment-specific duplicates within each cell type were collapsed by retaining the entry with the highest magnitude. Reporter scores are standardized within each parameterization and are therefore reported as medians indicating magnitude only. The same metabolites were scored in every parameterization, so ranks are compared directly.

Prostaglandin J2 (PGJ2) in the astrocyte was the highest-ranked reporter metabolite in 62 of 100 parameterizations (median reporter score 40.52; median rank 1, 5th–95th percentile 1–6), and astrocytic O-propanoylcarnitine ranked among the three highest-scoring entries in all 100. The remaining entries belonged to the linoleate pathway and comprised linolenic acid hydroperoxide, linoleyl carnitine, linoleoyl-CoA, and linoleate (Table 3 and Fig 6D). Ten cell-type-specific entries representing six unique metabolites are reported from 16 entries recovered across the 100 parameterizations, and every reported entry was suppressed in the knockout whenever it was scored. The complete reporter-metabolite output is provided in S11 Table.

**Table 3.**
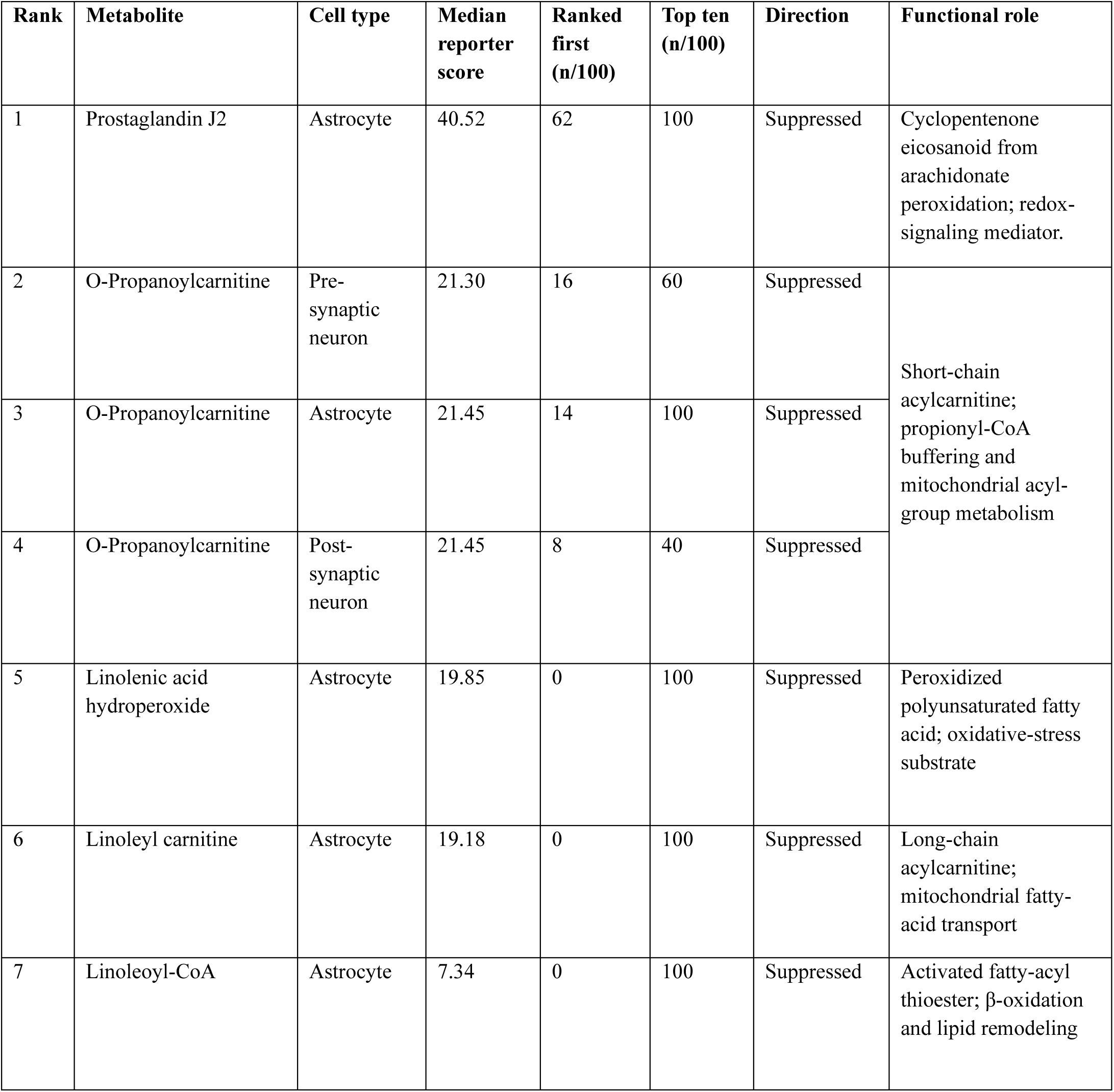
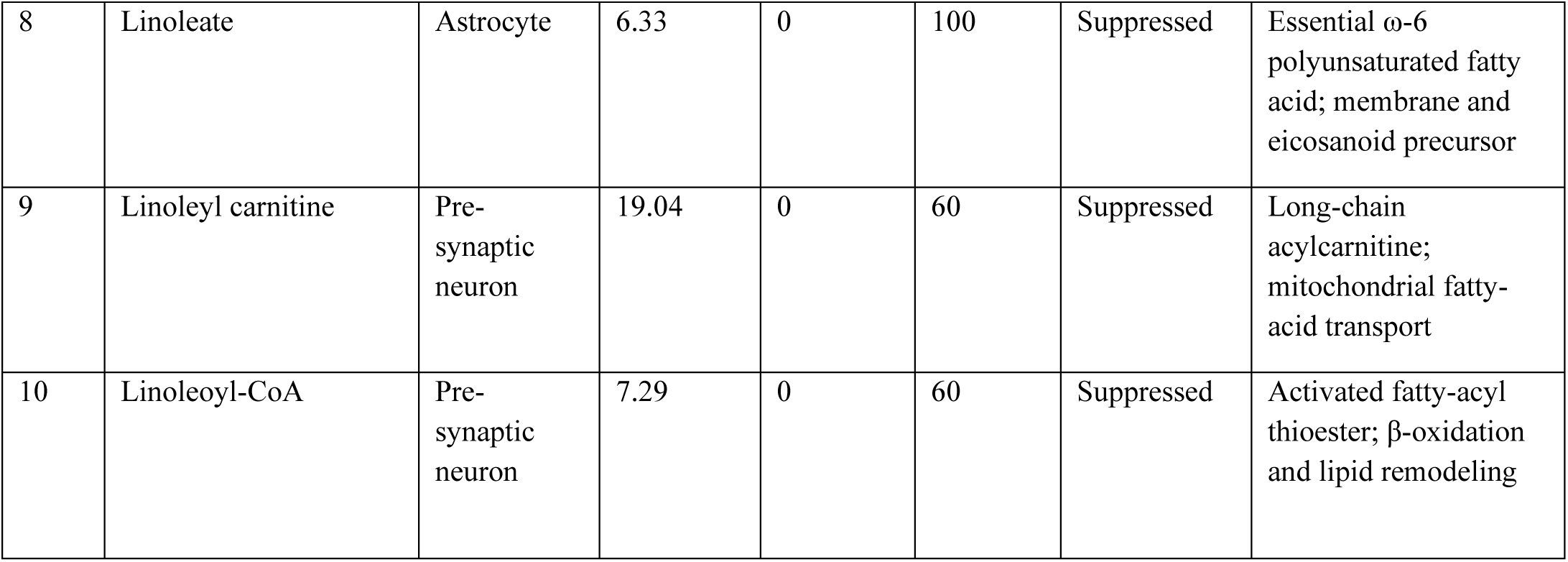
Lipid reporter metabolites altered in the FABP7 knockout under the ATP-maintenance objective across 100 turnover-number parameterizations. Entries are the ten highest-ranked metabolite–cell type combinations; the same metabolite may therefore appear in multiple cell types. In Ranked first (n/100) and Top ten (n/100), n denotes the number of parameterizations in which each entry ranked first or among the ten highest-scoring, respectively. Median reporter scores indicate magnitude only because scores are standardized within each parameterization; ranks are used for comparison across parameterizations. Direction indicates whether summed absolute flux decreased (suppressed) or increased (induced) in the knockout. Complete output is provided in S11 Table.

Prostaglandin J2 is a cyclopentenone eicosanoid derived from arachidonic acid. Cyclooxygenase converts arachidonate to prostaglandin D2, the most abundant prostaglandin in the brain, which subsequently undergoes non-enzymatic dehydration to prostaglandin J2 and its highly reactive metabolite 15-deoxy-Δ12,14-prostaglandin J2 [60]. Because prostaglandin J2 forms non-enzymatically from prostaglandin D2, which has been shown to suppress seizures and promote postictal sleep [61], the altered prostaglandin J2 flux predicted here occurs downstream of a prostaglandin pathway with established seizure-suppressive activity. The electrophilic cyclopentenone ring of prostaglandin J2 forms covalent adducts with glutathione and protein cysteine residues. As a result, prostaglandin J2 accumulates in the brain under neuroinflammation and oxidative stress and has been implicated in protein aggregation and neurodegeneration in Alzheimer’s and Parkinson’s disease [60, 62]. In the model, PGJ2’s suppression in astrocytes aligns with reduced routing of arachidonate-derived substrates into downstream eicosanoid signaling following disruption of astrocytic lipid trafficking, thereby linking FABP7-dependent lipid handling to glutathione-associated redox metabolism. The accompanying suppression of linoleate-pathway carnitine and coenzyme A esters indicates a coordinate reduction in astrocytic polyunsaturated-fatty-acid handling rather than an isolated eicosanoid effect.

By contrast, the biomass objective produced a less stable reporter ranking: prostaglandin J2 ranked first in 30 of 100 parameterizations, entered the ranking in 90, and was suppressed in the astrocyte in 45 of those, while the direction of change was inconsistent for most entries (Fig 5D and S11 Table).

## Discussion

In this work, we developed an enzyme-constrained genome-scale metabolic model of the tripartite synapse to investigate how proteome allocation shapes neuron–astrocyte metabolic coupling. The ecMMU1868-TPS model explicitly represents metabolite exchange across the synaptic cleft, protein capacity in each cell type (astrocyte, pre-synaptic neuron, and post-synaptic neuron), and FABP7-dependent lipid trafficking. This approach provides a quantitative framework for analyzing system-level metabolic interaction at the tripartite synapse (Fig 1 and 2). By incorporating enzymatic constraints into the cell-type-specific astrocyte and neuronal models and integrating them with a modified SteadyCom framework, this model addresses key questions in brain metabolism, including the conditions under which astrocyte-to-neuron lactate exchange becomes necessary and how lipid trafficking and redox buffering are coordinated across interacting brain cell types in the model [1].

The reconstructed models address two major gaps: the need for a high-quality generic mouse metabolic model and the lack of an enzyme-constrained tripartite synapse community model. Curation of initial mouse metabolic model, iMM1865, into iMMU1867 resolved 290 mass and charge imbalances, expanded the C4–C18 fatty-acid elongation and lipid pathways, and standardized gene–reaction and Systems Biology Ontology (SBO) annotations. These changes raised the MEMOTE [37] score from 46% to 83% (Fig 3A), corrected the physiologically unrealistic growth rate of 798 h⁻^1^ in iMM1865 to 0.327 h⁻^1^ [30], and achieved near-complete carbon closure (Table 1). Phenotypic testing and cell-type-specific reconstruction with iMAT [26] confirmed physiologically consistent, brain-relevant behavior and identified a conserved core of 299 reactions alongside astrocyte- and neuron-enriched pathways (Fig 3B, Fig 4 and Table 2).

Several mechanisms have been proposed to explain astrocyte–neuron lactate shuttle (ANLS), including activity-dependent glutamate signaling and differences in cellular energy demand and metabolic specialization [4, 54]. Our results do not exclude these possibilities. Instead, they show that enzyme-capacity limitations are sufficient to drive lactate shuttling in the tripartite synapse model, suggesting that proteome allocation may contribute to ANLS under high metabolic demand. Under ATP-maintenance, the primary objective here because mature brain cells are largely non-proliferative, the median lactate coupling onset across 100 turnover-number parameterizations was 0.770 of maximal demand, compared with 0.879 under biomass production, and the maintenance objective gave the earlier onset in every paired parameterization (Fig 5B and 6B). Thus, ANLS coupling reflects a broader response to enzyme-capacity constraints under different cellular objectives. Results from the stoichiometry-only tripartite synapse community model (iMMU1868-TPS) showed that the minimum feasible lactate exchange remained at zero across objective ranges, indicating that astrocyte-to-neuron lactate transfer was permissible but not obligatory coupled to either objectives (Fig 5A and 6A). Since iMMU1868-TPS and ecMMU1868-TPS share the same reaction network structure and differ only in proteome-allocation constraints, this lactate overflow is attributable to protein-capacity limitations rather than to network connectivity alone, mechanistically resembling proteome-constrained overflow metabolism observed in microbial systems [13, 14].

The proteome-allocation results from ecMMU1868-TPS provide a framework for integrating experimental observations on neuron–astrocyte metabolic coupling that emphasize either neuronal glycolysis or lactate-mediated support. *In vivo* imaging with a genetically encoded biosensor has revealed a lactate gradient from astrocytes to neurons, consistent with carrier-mediated transfer [63]. In contrast, activity imaging studies show that stimulated neurons engage in their own glycolysis and transiently increase the cytosolic NADH/NAD⁺ ratio rather than relying on lactate uptake [64]. Neurons also sustain aerobic glycolysis in their somata as an antioxidant mechanism, indicating that neuronal glucose metabolism is not merely a backup to astrocytic lactate supply [65]. Together, these observations suggest that neuronal glycolysis and lactate-mediated support may coexist across different metabolic states rather than represent competing models of brain energy metabolism. Within this framework, the experimental findings may reflect different operating points along the ATPM-maintenance- and biomass–lactate envelopes, with astrocytic lactate flux remaining permissive at lower demands and becoming obligatorily coupled above objective-specific thresholds. This interpretation aligns with the ANLS framework, in which neuronal glycolysis complements rather than replaces astrocyte-derived lactate support [1, 6].

The modeling framework identified FABP7 as a critical mediator of astrocytic lipid detoxification, establishing a connection between lipid trafficking and neuronal redox homeostasis. Simulating FABP7 deletion in ecMMU1868-TPS reduced astrocytic clearance capacity to 3.7% of wild-type, with the residual carried by FABP5 and FABP3 (Fig 5C and 6C). Because FABP7 is the dominant intracellular lipid-trafficking carrier supporting glutathione-dependent detoxification in astrocytes, its deletion reduced astrocytic processing of neuron-derived peroxidized lipids and increased neuronal oxidative burden. This prediction aligns with experimental evidence that peroxidized lipids generated in neurons during hyperexcitability are transferred to astrocytes via apolipoproteins (ApoE) for detoxification [12, 20] and that FABP7 further protects astrocytes from ROS-associated lipid toxicity by sequestering fatty acids into lipid droplets [19, 20].

In addition to its roles in lipid detoxification and redox balance, loss of FABP7 resulted in distinct, cell-type-specific changes in lipid metabolism. Reporter-metabolite analysis under ATP-maintenance objectives identified astrocytic prostaglandin J2 (PGJ2) as the highest-ranked metabolite in 62 of 100 turnover-number parameterizations and among the top ten in all 100, with consistent suppression in the knockout (Table 3 and Fig 6D). Other highly ranked metabolites included O-propanoylcarnitine and linoleate-associated metabolites, indicating coordinated changes in eicosanoid, acylcarnitine, and polyunsaturated-fatty-acid metabolism. PGJ2 is a cyclopentenone eicosanoid derived from arachidonic acid metabolism whose downstream metabolites accumulate during neuroinflammation and oxidative stress, contributing to protein aggregation and neurodegenerative processes through their highly reactive electrophilic cyclopentenone ring [60, 62]. PGJ2 forms non-enzymatically from prostaglandin D2, a prostaglandin implicated in seizure suppression and postictal sleep [61]. The predicted suppression of astrocytic PGJ2, together with changes in linoleate-pathway metabolites, therefore supports a broader disruption of FABP7-dependent lipid trafficking and redox-associated lipid metabolism. Under the biomass objective, reporter rankings and directions were less stable across parameterizations (Fig 5D and S11 Table), indicating that the detailed reporter-metabolite is objective-dependent.

These findings build upon previous genome-scale metabolic modeling efforts in brain metabolism. Earlier multicellular brain reconstructions showed that lactate exchange between brain cell types can be represented within stoichiometric network models constrained by measured cerebral metabolic rates [66], and that cell-type-specific astrocyte reconstructions can reproduce known metabolic capabilities [24, 30]. However, these models either lacked explicit neuron–astrocyte coupling or failed to provide a mechanistic explanation of the conditions under which lactate exchange is favored over alternative carbon-utilization routes [1–4]. By imposing enzyme-capacity constraints on a curated mouse model and integrating the resulting resource-allocation models into a tripartite synapse community model, this study shows that proteome limitation alone, without setting lactate exchange as an objective, is sufficient to produce obligatory astrocyte-to-neuron lactate flux under both ATP-maintenance and biomass objectives. The framework further links ANLS analysis to lipid peroxidation and to FABP7-dependent lipid trafficking and detoxification, enabling a coordinated investigation of metabolic support and lipid–redox regulation across the neuron–astrocyte interface within a single modeling framework. The model addresses several processes emphasized within the neuron–astrocyte metabolic unit concept [1], including astrocytic lactate support and detoxification of neuron-derived peroxidized lipids, and can be used to evaluate when these become limiting.

More broadly, the tripartite synapse framework provides a scalable platform for investigating metabolic dysfunction across neurological disorders, including epilepsy, Alzheimer’s disease, traumatic brain injury (TBI), and cerebral ischemia [7–10]. TBI is a particularly relevant application because FABP7 is strongly upregulated in reactive astrocytes following injury and is linked to both intracellular lipid handling and injury-responsive pathways that maintain blood– brain barrier integrity and regulate acetyl-CoA-dependent signaling [20, 67, 68]. Although these signaling mechanisms are not explicitly represented in the current model, ecMMU1868-TPS captures the underlying metabolic processes associated with FABP7 function, including ketone metabolism, lactate exchange, fatty-acid oxidation, lipid trafficking, and redox buffering. Epilepsy is another direct application, as the metabolic state strongly influences neuronal excitability. Recent experimental evidence indicates that FABP7-knockout mice exhibit elevated electroshock seizure thresholds during the dark (wake) phase, suggesting that FABP7 loss alters metabolic conditions associated with seizure susceptibility [69]. Since these metabolic processes are represented within ecMMU1868-TPS, the framework can be extended to examine how shifts in substrate utilization affect proteome allocation and metabolic coupling across the tripartite synapse. Future incorporation of condition-specific enzyme constraints derived from phase-resolved proteomics and circadian physiological parameters may further enable investigation of how time-of-day-dependent metabolism influences astrocyte–neuron coupling and FABP7-associated lipid and redox regulation [69].

While the present framework provides mechanistic insight into neuron–astrocyte metabolic coupling, several limitations should be considered when interpreting the results. First, the modeling framework operates under steady-state assumptions and does not capture the temporal dynamics of neuronal activity or time-dependent shifts in metabolic state. Activity-dependent transitions in the ANLS behavior can therefore be inferred only from changes in the feasible metabolic state relative to the protein-capacity threshold. Second, enzyme-constrained predictions depend on the accuracy and completeness of enzyme turnover number (*k*_cat_) assignments and the quality of experimentally measured proteomics data. In this study, *k*_cat_ values were assigned following the standard GECKO 3.0 protocol [15]. Although this strategy improves enzyme coverage and model feasibility, uncertainty in *k*_cat_ assignment may still influence predicted enzyme allocation and flux distributions. However, the bootstrap parameterization showed that the main lactate-coupling result was preserved across alternative *k*_cat_ assignments. Third, the reconstruction focuses on neuron–astrocyte metabolic coupling and may not capture metabolic variation across other brain cell types and regions. In addition, the model assumes a fixed tripartite synapse architecture and does not incorporate dynamic changes in perisynaptic astrocytic process (PAP) coverage, which can vary across brain regions, physiological states, and species. Incorporating dynamic PAP remodeling in future models may yield further insights into sleep, circadian rhythms, learning, and disease-related alterations in neuron–astrocyte interactions. Fourth, biomass production in mature, largely non-proliferative astrocytes and neurons should be interpreted as a proxy for macromolecular turnover and cellular homeostasis [30–32]. Fifth, the lipid-peroxidation simulations used a single peroxidized lipid species as the imposed oxidative challenge. Although sensitivity analysis was performed across varying input levels of this lipid, the model did not assess combinations or ratios of multiple peroxidized lipids, which may more accurately reflect diverse physiological and pathological states. Future research could investigate mixed lipid-peroxidation challenges to evaluate the robustness of predicted detoxification and redox responses. Sixth, the model does not explicitly represent pH dynamics or regulatory mechanisms governing substrate utilization. While proton stoichiometry and enzyme-capacity constraints are incorporated, the model omits metabolite concentrations, buffering effects, pH-dependent kinetics, and regulation of substrate preference. These acid-base interactions may become particularly significant under ketogenic conditions and will require explicit modeling in future studies. Seventh, FABP transport was parameterized using DLKcat-predicted turnover numbers, despite FABPs being non-catalytic lipid-binding proteins and thus outside the predictor’s training domain. The 3.7% residual is contingent upon the prediction that FABP5’s turnover is 24.2 times lower than that of FABP7; therefore, this value should be interpreted as conditional on the DLKcat predictions. Eighth, SteadyCom constrains the three cell models to a common growth rate, thereby fixing their relative abundances. Given that proteome allocation and maintenance demands vary among cell types, the onset of lactate coupling may depend on the specific community composition. Finally, the cell-type-specific reporter-metabolite redistribution (Table 3) predicted following FABP7 loss remains to be experimentally validated.

In summary, this study demonstrates that enzyme-capacity constraints are sufficient to drive lactate shuttling, in which astrocyte-derived lactate becomes increasingly required for neuronal metabolism across both ATP-maintenance and biomass-production objectives. The model also shows that FABP7-dependent lipid trafficking links astrocytic detoxification with neuronal redox balance, connecting lactate shuttling and lipid–redox handling within a single proteome-constrained tripartite synapse framework. The cell-type-specific reporter-metabolite signature predicted for FABP7 loss underscores how disruption of lipid trafficking can redistribute lipid-associated flux across astrocytic, presynaptic, and postsynaptic compartments. Furthermore, the enzyme-constrained tripartite synapse model developed in this study serves as a cellular-scale building block for a planned multi-scale gut–liver–brain modeling framework. Beyond these findings, the model’s explicit representation of fatty acid and lipid metabolism offers a framework for addressing unresolved questions in brain metabolism and for identifying metabolic vulnerabilities across various neurological disease states, such as circadian and ketogenic regulation in epilepsy and lipid–redox alterations following traumatic brain injury [20, 67, 70].

## Materials and Methods

The computational workflow for reconstructing and analyzing the tripartite synapse model consisted of five main stages (Fig 1B): (i) evaluation and curation of the generic mouse metabolic model iMM1865; (ii) derivation of astrocyte- and neuron-specific models using iMAT; (iii) expansion to a tripartite synapse architecture; (iv) reconstruction of enzyme-constrained resource allocation models with GECKO 3.0; and (v) multicellular coupling using a modified SteadyCom framework. Each stage is described in detail below.

### Evaluation of the mouse metabolic model, iMM1865

The evaluation of the iMM1865 [25] mouse genome-scale metabolic model began by generating a MEMOTE [37] report to assess consistency with current model quality standards. The MEMOTE report identified deficiencies in mass and charge balance (290 imbalances) and annotation coverage, with an overall score of 46% (Fig 3A). Flux balance analysis (FBA) and flux variability analysis (FVA) were then performed to evaluate biomass production (yielding an unrealistic value of 798 h⁻¹; Table 1) and reaction flux distributions. All detected inconsistencies were systematically addressed through manual curation. The original and updated MEMOTE reports, as well as the curated iMMU1867 model files, are available in the GitHub repository associated with this study (https://github.com/SchroederLabWSU/ecMMU1868-TPS).

### iMM1865 model curation

The iMM1865 model served as the base genome-scale metabolic reconstruction, comprising 10,612 reactions, 5,837 metabolites, and 1,865 genes. Curation addressed four categories of deficiencies:

**(1) Exchange reaction correction**We first identified exchange reactions directly from the SBML file using regular-expression searches for exchange-reaction identifiers, which confirmed that every exchange reaction in the distributed model carried the default lower bound of −1000 mmol·gDW⁻^1^·h⁻^1^. Carbon balance analysis then quantified and compared total carbon uptake with total carbon secretion (including biomass production) under parsimonious flux balance analysis (pFBA) to verify elemental conservation. Exchange reactions acting as major unintended carbon sources and sinks were blocked, uptake was restricted to the curated medium, and the carbon balance was recomputed. This was repeated until the carbon output distribution was dominated by physiologically expected species, such as lactate, carbon dioxide, and bicarbonate.
**(2) Mass and charge balancing.** All 290 mass and charge imbalances were resolved by correcting molecular formulas, charges, and reaction stoichiometry. Imbalanced reactions were identified using MEMOTE and a Jupyter notebook script based on the check_mass_balance function from COBRApy [71]. Key metabolite corrections, which resolved numerous reaction imbalances, included orthophosphate (M_pi; charge corrected from 0 to −2), adenosine 3,5-bismonophosphate (M_pap; charge corrected from 0 to −4), and dopamine (M_dopa; molecular formula corrected from C_8_H_11_NO_2_ to C_8_H_12_NO_2_). Corrections were performed using biochemical databases, including ModelSEED [72], BiGG [73], and KEGG [29]. Another major source of imbalance was the use of undefined acyl R-groups in phospholipid reactions. To address this, a fatty acid pool abstraction was introduced, collapsing individual acyl-resolved phosphatidylcholine species into an averaged phosphatidylcholine pool. Brain fatty acids are a distribution of species rather than a single compound, so the pool metabolite was assigned the abundance-weighted average composition of the 15 fatty acid species reported for mouse brain tissue [36]. The weighted average is C_17.7727_ H_31.7184_O_2_ with charge −1, which is not an integer composition. Because SBML chemical formulas require integer subscripts, the composition was scaled by 10,000, giving the fatty acid pool metabolite (M_Fattotal_c) the molecular formula C_177727_H_317184_O_20000_ with charge of −10000. Stoichiometric coefficients were correspondingly divided by 10,000, so that one unit of flux still corresponds to one average fatty acid molecule. The formula scaling preserves the integer SBML formula, while the coefficient scaling normalizes the reaction to a single pool unit, together preserving elemental mass balance. Acyl-resolved phosphatidylcholine reactions containing the “R” group element in metabolite formulas were removed only after confirming that their metabolic conversions were retained by the corresponding pool-based reactions. The curated model was then tested for growth on the defined medium, and mass and charge balance were recalculated, confirming that the pool-based formulation improved elemental consistency.
**(3) Fatty acid and lipid pathway expansion.** Missing acyl-carrier protein (ACP)-associated reactions required for fatty acid biosynthesis initiation were reconstructed from the KEGG fatty acid biosynthesis pathway (map00061) [29]. Fatty acid and lipid pathways were further expanded to include reactions involved in lipid trafficking and peroxidation, such as linolenic acid peroxidation, glutathione-dependent lipid hydroperoxide reduction, and lipid alcohol repair pathways.
**(4) Model standardization and validation.** Missing reactions and metabolite annotations were identified by inspecting reactions flagged as mass- or charge-imbalanced by MEMOTE and COBRApy-based balance checks, tracing these imbalances to missing reactants, products, or annotations, and comparing the affected pathways against biochemical pathway databases including KEGG, ModelSEED, and BiGG. Systems Biology Ontology (SBO) terms were added to all reactions and species. The curated model quality and consistency were checked using MEMOTE. Four phenotypic validation tests were performed to examine model behavior: (i) growth response to a carbon-supply sweep for five key carbon sources (glucose, lactate, glutamine, palmitate, β-hydroxybutyrate); (ii) respiratory quotient (RQ) analysis of complete substrate oxidation for the same five carbon sources; (iii) oxygen sensitivity analysis by gradually restricting oxygen uptake; and (iv) a single-nutrient omission screen to identify essential nutrients. Degree of reduction was calculated per carbon as γ = (4n_C_ + n_H_ − 2n_O_ − 3n_N_)/n_C_, where n_C_, n_H_, n_O_, and n_N_ are the numbers of carbon, hydrogen, oxygen, and nitrogen atoms, respectively, using the neutral form of each substrate [38]. The complete validation Jupyter notebook is available in the project GitHub repository (https://github.com/SchroederLabWSU/ecMMU1868-TPS).

### Cell-specific reconstruction with iMAT

Astrocyte-specific (iMMU1867a) and neuron-specific (iMMU1867n) models were derived from the curated iMMU1867 model using iMAT [26], implemented in Python via the iMATpy package [74]. Cell–type–resolved proteomic data from a mouse brain dataset [27] served as the basis for reconstructing cell–type–specific models. The dataset included label-free quantification (LFQ) intensity values measured across cell type–enriched fractions of mouse brain tissue. The processed proteomics data used for iMAT implementation, including gene names and corresponding model identifiers, are provided in S2 Data.

**(1) Gene identifier mapping.** Gene identifiers in the model (Entrez Gene IDs) were mapped to their corresponding Entrez gene names. The proteomics dataset reported gene names, which were converted to Entrez Gene IDs and matched to the model gene identifiers using a custom Python script, solely for mapping proteomics data to the model for subsequent iMAT analysis.
**(2) Proteomics processing.** For each cell type, LFQ intensity values across three biological replicates were aggregated by computing the median. The resulting gene-level median expression values were used for subsequent gene weight assignment (S2 Data).
**(3) Gene weight assignment.** Gene expression values were discretized into qualitative weights following the standard iMAT protocol [74]. For each cell type, only genes with detected (non-zero median) expression were considered. Percentile thresholds were calculated from the distribution of detected gene expression values. Genes with expression above the upper percentile (top 15%) threshold were assigned a weight of +1 (high expression), those below the lower percentile (bottom 15%) threshold were assigned −1 (low expression), and all remaining genes were assigned 0 (moderate/unconstrained).
**(4) Reaction weight propagation.** Gene weights were propagated to reaction-level weights using the model’s GPR rules via the gene_to_rxn_weights function, which applies AND/OR logic consistent with the gene–protein–reaction mapping.
**(5) iMAT optimization.** The iMAT algorithm was run with epsilon set to 1×10⁻^3^ and threshold set to 1×10⁻^6^. Epsilon specifies the minimum flux for reactions associated with highly expressed genes to be considered active, while the threshold defines the maximum allowable flux for reactions associated with lowly expressed genes. Epsilon was selected based on flux distribution obtained using pFBA for the curated iMMU1867 model before iMAT was applied. The 387 active reactions separated into two distinct flux groups, with 348 carrying flux above 1 × 10⁻³ mmol·gDW⁻^1^·h⁻^1^ and the remaining 39 below 1 × 10⁻⁶ mmol·gDW⁻^1^·h⁻^1^. The median nonzero flux was 0.048 mmol·gDW⁻^1^·h⁻^1^. Therefore, epsilon at 1 × 10⁻³ lies below the magnitude of typical active fluxes while remaining clearly separated from very small flux values. The threshold value of 1 × 10⁻⁶ mmol·gDW⁻^1^·h⁻^1^ matches the iMAT off-threshold used to constrain low-expression reactions. It is also one order of magnitude above the solver tolerance of 1 × 10⁻⁷, ensuring the flux threshold remains above the solver’s numerical tolerance. The astrocyte- and neuron-specific context models were generated using the “subset” method in the iMATpy model_creation module, which preserves the full network topology and adjusts reaction bounds to reflect cell–type–specific metabolic activity patterns.

### Active reaction overlap analysis

Parsimonious flux balance analysis (pFBA) was performed on the generic (iMMU1867), astrocyte-specific (iMMU1867a), and neuron-specific (iMMU1867n) models using COBRApy to obtain parsimonious flux distributions that minimize total network flux while maintaining optimal biomass production. A reaction was classified as active if its absolute flux exceeded 1×10⁻^6^ mmol·gDW⁻^1^·h⁻^1^

### Extension to the tripartite synapse

The cell-type-specific models from iMAT were further expanded to represent the tripartite synapse architecture. iMMU1867a was extended with a perisynaptic astrocytic process (PAP) compartment (pap) to represent the fine astrocytic processes that ensheathe synapses (Fig 2). Two copies of the iMMU1867n were generated: one representing the pre-synaptic neuron (iMMU1868n-pre with a pre-synaptic compartment npre_c) and one representing the post-synaptic neuron (iMMU1868n-post, with a post-synaptic compartment npost_c). A shared synaptic cleft compartment (syn_e) was introduced to enable metabolite exchange between cells at the synapse. A lipid peroxidation module was added to represent the transfer of peroxidized lipid (C04717) from neurons to astrocytes for detoxification. The complete set of transport reactions, associated metabolites, and GPR associations was curated and manually added to the SBML files using Notepad++ and is detailed in S1 Data.

Key metabolic processes added during the tripartite expansion included H⁺-coupled MCT-type lactate transport between the astrocyte PAP and synaptic cleft, EAAT-type glutamate uptake into the astrocyte PAP with SNAT-type glutamine release, D/L-serine shuttle reactions, the GSH/Cys-Gly cycle for cross-cellular antioxidant support, and a lipid peroxidation and FABP7-dependent detoxification pathway (S1 Data).

### Enzyme-constrained model reconstruction

Enzyme-constrained astrocyte and neuron models were reconstructed using the GECKO 3.0 framework [15] with cell type–specific proteomics data [27]. Reconstruction was performed in MATLAB (R2025a) using the RAVEN Toolbox 2.10.4 and GECKO Toolbox 3.0, with Gurobi optimizer 12.0.2 serving as the linear programming (LP) solver.

**(1) Model import and adapter configuration.** For each cell type, a custom ecModelGEMAdapter was configured with organism-specific parameters (*Mus musculus*, KEGG: mmu), including total protein content (P_tot_ = 0.5 g protein·gDW⁻^1^), active enzyme fraction (σ = 0.5), and enzyme mass fraction (f = 0.5 g enzyme/g protein). These initial adapter values are GECKO defaults; the f-factor was subsequently recalculated from proteomics data using calculateFfactor, yielding cell type–specific values of f = 0.0278 for astrocytes and f = 0.0120 for both pre- and post-synaptic neurons. The biomass reaction (BIOMASS_reaction) and glucose exchange reaction (EX_glc D_e) were specified. Adapter files and parameter configurations are available in the GitHub repository associated with this study (https://github.com/SchroederLabWSU/ecMMU1868-TPS).
**(2) Enzyme-constrained model creation.** Enzyme-constrained models were generated using makeEcModel, which splits reversible reactions into forward and reverse reactions and introduces enzyme usage pseudometabolites and pseudoreactions. Models were created for pre-synaptic neurons (ecMMU1868n-pre), post-synaptic neurons (ecMMU1868n-post), and astrocyte (ecMMU1868a-pap) cell lines. Enzyme turnover numbers (*k*_cat_) were assigned using the standard GECKO turnover-number assignment workflow. EC annotations in the model were first used to retrieve experimentally reported turnover numbers from BRENDA. When an exact EC-number match was available, the corresponding BRENDA *k*_cat_ value was used. For reactions without an exact EC-matched BRENDA value, turnover numbers were assigned using DLKcat machine-learning predictions. EC-wildcard fuzzy matching was used only when neither an exact BRENDA match nor a DLKcat prediction was available, and the highest *k*_cat_ value was selected only within that fuzzy-matched category.
**(3) Protein pool constraint.** Protein pool constraints were applied using setProtPoolSize, with the fraction of measured protein (f-factor) computed via calculateFfactor and combined with P_tot_ and σ. When enzyme-capacity constraints reduced the maximum growth rate below that of the corresponding stoichiometric model, GECKO’s sensitivityTuning procedure identified the *k*_cat_ constraints that most strongly limited growth and iteratively adjusted them until the reference growth rate was recovered. This is part of the standard GECKO workflow rather than a manual adjustment of individual *k*_cat_ values. The procedure converged in 9 iterations and altered 6 of the 18,646 *k*_cat_ assignments in the model. The tuned reactions, enzymes, and original and adjusted *k*_cat_ values are listed in S3 Data.

Metabolic fluxes in the enzyme-constrained model are subject to the kinetic constraint:

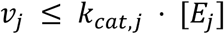

where v_j_ is the flux of reaction j, *k*_cat,j_ is the corresponding turnover number, and [E_j_] is the enzyme concentration. Total enzyme usage is bounded by:

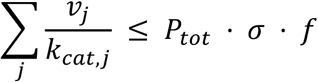

where Pₜₒₜ is total cellular protein, σ is the fraction of protein that is catalytically active, and f is the fraction of the proteome represented in the model.

### Tripartite synapse coupling using a modified SteadyCom framework

The enzyme-constrained astrocyte (ecMMU1868a-pap) and neuron models (ecMMU1868n-pre and ecMMU1868n-post) were combined using a modified SteadyCom workflow [28] to generate the enzyme-constrained tripartite community model (ecMMU1868-TPS) and simulate intercellular metabolic exchange at the tripartite synapse. The same procedure was applied to the stoichiometric astrocyte (iMMU1868a-pap) and neuron models (iMMU1868n-pre and iMMU1868n-post) to generate the stoichiometric-only tripartite synapse model (iMMU1868-TPS). We refer to the workflow as “modified” because the standard SteadyCom was originally formulated for stoichiometric community models, but here it was applied to enzyme-constrained models, in which each cell carries its own protein-pool constraint. In addition, a shared synaptic-cleft exchange pool (compartment [q]) was introduced alongside the shared extracellular pool ([u]) used in standard SteadyCom. The shared growth rate is the standard SteadyCom coupling variable used to enforce a balanced community state. In this study, we used it as a mathematical coupling condition for the aggregate macromolecular turnover and homeostatic demands represented by the biomass reactions across the three cell types. Each cell type still retains its own reaction network, flux distribution, and enzyme-capacity constraint, while the models are coupled through shared metabolite pools and the SteadyCom community constraints. SteadyCom was implemented in MATLAB using the COBRA Toolbox.

**(1) Community model construction.** The three enzyme-constrained models (ecMMU1868a-pap, ecMMU1868n-pre, ecMMU1868n-post) were loaded, and their metabolite identifiers were standardized from the metabolite_compartment format to the metabolite[compartment] format. The createMultipleSpeciesModel function was used to build a three-species community model with species tags AST, PRE, and POST. A shared extracellular pool (compartment [u]) was generated automatically by the community builder.
**(2) Synaptic cleft shared pool.** In addition to the standard shared extracellular pool, a second shared pool representing community exchange at the synaptic cleft was introduced to model synapse-specific metabolite interactions and separate them from general extracellular exchange. Metabolite exchange allowed through the synaptic cleft includes lactate, glutamate, glutamine, D-serine, L-serine, glycine, K⁺, Na⁺, H⁺, and lipid peroxidation intermediates.
**(3) Growth coupling.** Community growth was constrained such that all three cell types share a common growth rate, consistent with the SteadyCom assumption of a steady-state community at fixed relative abundances. This was implemented by coupling the three cell-specific biomass reactions to equal fluxes (v_AST = v_PRE = v_POST) through two pseudo-reactions, couple_AST_PRE and couple_AST_POST. For the enzyme-constrained community, each cell-specific protein-pool exchange was assigned a lower bound of −10 mmol·gDW⁻^1^·h⁻^1^. This value was selected from a sensitivity analysis of the prot_pool_exchange bound (−1000 to −0.001 mmol·gDW⁻^1^·h⁻^1^), which showed that maximum community growth remained unchanged at 0.105 h⁻¹ for capacities of 20 and above, fell progressively below that, and became infeasible below 0.02. The selected bound therefore lies within the range in which the GECKO-imposed enzyme-capacity constraints remain growth-limiting (S9 Table).
**(4) Medium constraints.** A defined minimal medium was designed to approximate brain interstitial fluid under glucose-fed conditions (S2 Table). D-Glucose was used as the primary carbon source with an uptake rate of 10 mmol·gDW⁻^1^·h⁻^1^ (60 C-mmol·gDW⁻^1^·h⁻^1^). Nine essential amino acids (EAAs) were supplied, reflecting the constraint that mammalian cells cannot synthesize EAAs *de novo* and must import them at physiologically limited rates. Individual uptake rates were determined by FVA at 99% of maximum biomass flux. For each EAA exchange reaction, FVA identified the minimum uptake rate required to sustain near-optimal growth, and the resulting bound was multiplied by a 5% buffer factor (×1.05) to prevent numerical infeasibility at the constraint boundary. This ensured that EAA uptake rates reflected the minimum physiologically necessary import, eliminating non-physiological overflow secretions observed under unconstrained medium conditions. Inorganic nutrients (ammonium, phosphate, sulfate) and oxygen were each allowed a maximum uptake of 10 mmol·gDW⁻^1^·h⁻^1^. Water, protons, and major ions (Na⁺, K⁺, Cl⁻, Ca²⁺) were left unconstrained (set arbitrarily to −1000 to 1000 mmol·gDW⁻^1^·h⁻^1^). Because these uptake bounds were selected to reproduce the astrocyte biomass range reported by Angarita-Rodriguez et al. [30], the resulting iMMU1867 biomass flux of 0.327 h⁻¹ is treated as a fitted reference value rather than an independent prediction. Uptake bounds for the SteadyCom community model were mapped from individual species exchange reactions to community exchange reactions (EXcom), preserving the same medium composition.

### Lactate coupling envelope analysis

Biomass–lactate production envelopes were computed for the stoichiometric tripartite synapse community model (iMMU1868-TPS) and the enzyme-constrained tripartite synapse community model (ecMMU1868-TPS). The same defined minimal medium used for the curate model, iMMU1867, was applied to the SteadyCom community model at the shared extracellular compartment, denoted u in the model. All exchange reactions into this shared pool were initially closed, and only the defined medium metabolites were reopened. This approach ensured that nutrients entered the three-cell community through a single shared extracellular pool, with uptake constrained at the community level rather than separately for each cell type.

Lactate coupling was evaluated under both ATPM and biomass objectives. For the ATP-maintenance analysis, biomass production was fixed at zero in all three cell types, and the objective was defined as the sum of fluxes through AST_ATPM, PRE_ATPM, and POST_ATPM. The maximum feasible ATP-maintenance flux was determined, and ATPM– lactate envelopes were generated at 64 evenly spaced fractions of this maximum by minimizing or maximizing EX_lac L[u]. For the biomass analysis, the maximum feasible community growth rate (µ_max_) was determined, and biomass–lactate envelopes were generated by fixing the coupled biomass flux at 64 evenly spaced fractions of µ_max_ from 0 to 1 and minimizing or maximizing EX_lac L[u].

To assess the dependence of lactate coupling on turnover-number assignment,100 *k*_cat_ bootstrap parameterizations of ecMMU1868-TPS were performed. For each parameterization, every reaction with both a BRENDA value and a DLKcat prediction was independently assigned one of the two available *k*_cat_ values, while reactions with a single estimate retained their values. EC-wildcard fuzzy-matched values were also kept fixed, as they were used only when neither a BRENDA nor a DLKcat estimate was available. Among the 11,786 reactions with paired estimates, 11,582 are present in ecMMU1868-TPS and were resampled. BRENDA exceeded DLKcat in 76% of cases by a median factor of 16. For each resulting parameterization, the lactate-coupling onset was recomputed as the lowest growth fraction at which minimum feasible lactate secretion exceeded 1% of the maximum feasible secretion at the same growth rate, with the onset located using a bisection root-finding method. The 1% criterion was used as a numerical threshold rather than a biological cutoff. We used the same 100 *k*_cat_ parameterizations for the biomass-production and ATP-maintenance analyses, allowing paired comparison of coupling onset between the two objectives.

### ROS and lipid peroxidation simulations

A ROS generation and lipid peroxidation module [75] was integrated into the tripartite synapse framework. All simulations were performed using the enzyme-constrained tripartite synapse community model, ecMMU1868-TPS. Peroxidized linoleate was represented as 13-L-hydroperoxylinoleic acid (13-HPODE; C04717). FABP7-mediated transport was modeled by AST_C04717tc (Entrez Gene ID: 12140), while parallel FABP5- and FABP3-mediated transport reactions, AST_C04717tcFABP5_EXP_1 (Entrez Gene ID: 16592) and AST_C04717tcFABP5_EXP_2 (Entrez Gene ID: 14077), were included as alternative C04717 carriers. FABP5 and FABP3 were selected due to their expression as brain fatty acid-binding proteins [76, 77], their gene representation in the reconstruction, and their detection in the astrocyte proteomics dataset [27]. DLKcat-predicted *k*_cat_ values for C04717 transport were 6.2826 s⁻¹ for FABP7, 0.2594 s⁻¹ for FABP5, and 3.8233 s⁻¹ for FABP3. FABP7 accounted for an astrocytic proteome share of 5.820 × 10^−4^, while FABP5 and FABP3 abundances were 0.5393 and 0.0074 times that of FABP7, respectively. Multiplying these values by the astrocyte protein-pool bound of 10 mmol·gDW⁻¹·h⁻¹ yielded protein-usage capacities of 5.820 × 10⁻³, 3.139 × 10⁻³ and 4.307 × 10⁻⁵ mmol·gDW⁻¹·h⁻¹ for FABP7, FABP5 and FABP3, respectively. The maximum transport capacity for each carrier was calculated by dividing its protein-usage cap by the corresponding enzyme stoichiometric coefficient (0.9115, 15.4363, and 1.0767), resulting in 6.385 × 10⁻³, 2.033 × 10⁻⁴, and 4.000 × 10⁻⁵ mmol·gDW⁻¹·h⁻¹ (S10 Table).

Lipid-peroxidation analysis was conducted under both biomass-production and ATPM objectives. For ATPM, the community biomass reaction was fixed at zero, and the summed ATPM flux across the three cell types was maximized. C04717 was supplied via PRE_SK_C04717_prec, and clearance was quantified as the total flux through downstream glutathione-dependent reduction reactions. The FABP7-knockout model was generated by deleting the FABP7 gene. FVA was performed with the community biomass objective fixed at its optimum for both genotypes to determine the feasible range of C04717 clearance across alternative optimal solutions. To assess knockout sensitivity to the FABP5 and FABP3 *k*_cat_ values, these values were scaled by 0.1, 0.32, 1, 3.2, and 10, spanning ±1 log₁₀ unit around the DLKcat predictions, consistent with the reported prediction error of DLKcat [58].

### Reporter metabolite analysis

The reporter score quantifies the extent to which reactions surrounding a metabolite change, scaled by the number of reactions connected to it. Reporter metabolite scores were computed from absolute flux differences between WT and FABP7 knockout simulations under both the biomass-production and ATP-maintenance objectives, following the method of Patil and Nielsen [59].

For each reaction *j*, the flux difference was calculated as:

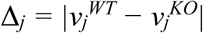

where v_j_^WT^ and v_j_^KO^ are the fluxes of reaction j in the wild-type and FABP7-knockout simulations, respectively. These values were standardized into *z*-scores across all active reactions:

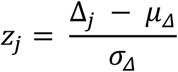

where μ_Δ_ and σ_Δ_ are the mean and standard deviation, respectively, of the flux-difference distribution Δj, calculated over reactions active in either the wild-type or knockout simulation.

For each metabolite *i*, the reporter score was computed as:

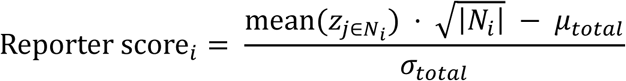

where *N_i_* is the set of reactions connected to metabolite *i*, |*N_i_*| is the number of such reactions, and μ_total_ and σ_total_ are the mean and standard deviation of the distribution of aggregated scores across all metabolites. Metabolites were ranked by the absolute value of the reporter score. High absolute scores identify metabolite nodes around which coordinated flux redistribution is concentrated. The analysis was implemented in Python, and the complete reporter metabolite table is provided as S11 Table.

For directionality, the signed flux-change score for each metabolite was computed separately as:

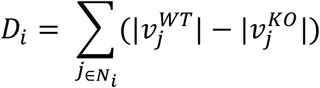

Where D_i_ is the directional score for the metabolite *i*. Metabolites with D_i_ > 0 were classified as suppressed, indicating lower summed absolute flux in FABP7 KO relative to WT. Metabolites with D_i_ < 0 were classified as induced, indicating higher summed absolute flux in the FABP7 KO relative to WT.

## Supporting information

**S1 Fig. Lactate-coupling onset across biomass and ATPM objectives.** Histogram comparing lactate-coupling onset distributions under the biomass and ATPM objectives across the same 100 turnover-number bootstrap parameterizations of ecMMU1868-TPS. Data are provided in S8 Table. (TIF)

**S2 Fig. Sensitivity of FABP7-knockout peroxidized-lipid clearance to FABP5 and FABP3 turnover numbers.** Astrocytic clearance of neuron-derived peroxidized lipid (C04717) against delivered load for WT and FABP7-KO at the DLKcat-predicted turnover numbers and for FABP7-KO with FABP5 and FABP3 *k*_cat_ values scaled across ±1 log₁₀ of their predictions. (TIF)

**S1 Data. Intercellular transport and lipid peroxidation reactions.**

(XML)

**S2 Data. Processed astrocyte and neuron cell-type-specific proteomics data.**

(CSV)

**S3 Data. Turnover numbers adjusted by GECKO sensitivity tuning in ecMMU1868-TPS.**

(CSV)

## Author Contributions

**Conceptualization:** Samuel Hayford Ayensu, Carlos C. Flores, Jason R. Gerstner, Wheaton L. Schroeder.

**Data curation:** Samuel Hayford Ayensu.

**Formal analysis:** Samuel Hayford Ayensu.

**Funding acquisition:** Wheaton L. Schroeder.

**Investigation:** Samuel Hayford Ayensu.

**Methodology:** Samuel Hayford Ayensu, Wheaton L. Schroeder.

**Project administration:** Wheaton L. Schroeder.

**Software:** Samuel Hayford Ayensu, Dan Hideki Horimoto.

**Supervision:** Wheaton L. Schroeder.

**Validation:** Samuel Hayford Ayensu, Dan Hideki Horimoto.

**Visualization:** Samuel Hayford Ayensu.

**Writing – original draft:** Samuel Hayford Ayensu.

**Writing – review & editing:** Samuel Hayford Ayensu, Dan Hideki Horimoto, Carlos C. Flores, Jason R. Gerstner, Wheaton L. Schroeder.

## Acknowledgements

This research used the resources of the Center for Institutional Research Computing at Washington State University.

## Data availability statement

All data, codes, and supplementary files are available on a GitHub repository at https://github.com/SchroederLabWSU/ecMMU1868-TPS. The large model files that exceed GitHub’s file size limit are archived on Zenodo and available at: https://doi.org/10.5281/zenodo.23225222

